# Remote sensing reveals multi-decadal losses of tree cover in California driven by increasing fire disturbance and climate stress

**DOI:** 10.1101/2021.11.30.470651

**Authors:** Jonathan A. Wang, James T. Randerson, Michael L. Goulden, Clarke Knight, John B. Battles

## Abstract

Forests provide natural climate solutions for sequestering carbon and mitigating climate change yet are threatened by increasing temperatures and disturbance. Accurate information on vegetation dynamics is lacking in some regions with forest carbon offset programs and dense forests like California. To address this, we combined remote sensing observations with geospatial databases to develop annual maps of vegetation cover (tree, shrub, herbaceous) and disturbance type (fires, harvest, and forest die-off) in California at 30 m resolution from 1985 to 2021. California lost 3783 km^2^ of its tree cover area (5.5% relative to initial cover). Early gains in tree cover area were more than offset by fire-driven declines, resulting in greater shrub and herbaceous cover area. Fires and tree cover area loss occurred where temperatures were high or increasing, whereas tree cover gain occurred in cooler areas. Disturbance and warming are threatening the integrity of California’s forests and its carbon offsets program.

**Teaser:** Climate and disturbance-driven tree cover loss challenges the viability of forests as natural climate solutions in California

## Introduction

Land management policies designed to reduce greenhouse gas emissions can provide much of the climate mitigation needed to stabilize warming below 2° C (*1*). In the United States, natural climate solutions could account for as much as 21% of net annual emissions; forests contribute the vast majority of this climate mitigation potential (*2*). In California, forest management aimed at reducing these threats are a focus of the state’s efforts to reduce wildfire hazard and improve forest resilience (*3*) but their efficacy is hampered by a lack of robust quantification of California’s forest resources. California’s legislature has mandated reductions in greenhouse gas emissions (*4, 5*), yet estimates of carbon dynamics in California’s forests remains highly uncertain with some studies reporting losses (*6*) and some reporting gains (*7*). It is therefore crucial to improve understanding of the distribution and dynamics of vegetation cover and their provisioning of ecosystem services like carbon sequestration and wildfire risk (*8*).

However, anthropogenic climate change and human activity are threatening the integrity of forests throughout the American West, and in California in particular (*9*), which may threaten the viability of natural and working lands as natural climate solutions (*10–12*). Temperate ecosystems such as those in California are diverse, dynamic, and climate-sensitive (*13*). As a result, climate change is driving substantial recent changes in ecosystem composition and structure (*14*), including warming-driven upward vegetation shifts in the mountains (*15, 16*), drought-induced mortality in the Sierra Nevada (*17, 18*), and increasing wildfire impacts throughout the forested mountains (*19, 20*). These changes in ecosystem structure and composition increasingly challenges the capacity of natural climate solutions to mitigate the impacts of anthropogenic fossil fuel emissions (*12*).

Forest management policies attempt to reduce forest vulnerability by reducing the density of fuels by direct removal (*21–23*) or prescribed burns (*24*). These changes in vegetation cover are intended to reduce sensitivity to drought and fire (*10, 25*). However, efficient implementation of these practices is limited by the considerable uncertainty in the highly spatially heterogeneous dynamics of fires (*26, 27*), timber harvest (*28*), and forest die-off due to drought, pests, and pathogens (*29, 30*). Furthermore, there is little quantification of the potential interactions of these different disturbance agents, though studies increasingly recognize the potential for pests and pathogens to increase fire susceptibility (*31*). Relying upon sparse field plots, such as the US Forest Service’s Forest Inventory and Analysis dataset, may result in significant variance and potential biases in the estimation of carbon stocks and vegetation cover (*32*). Timely geospatial datasets of vegetation cover and other ecosystem properties are thus essential to comprehensively characterize vegetation change, reduce bias in ecosystem monitoring, and efficiently manage forests to meet climate action goals.

Individual studies and datasets have quantified changes in forest dynamics due to drought-driven die-off (*17, 33*), fires (*34*), and management (*28*), but these studies typically consider only individual disturbance agents and span relatively short time-scales (*35*). Existing geospatial datasets of disturbance often are spatially coarse and assume spatial homogeneity within disturbance perimeters, despite the frequent occurrence, for example, of unburned islands that can occur frequently in forest fires (*26, 36*). There remains a need to compare how these different disturbance agents are reshaping tree cover in managed ecosystems like in California on multi-decadal scales in order to better quantify the carbon balance and other ecosystem services (*37*), particularly as climate change alters disturbance regimes (*38, 39*).

California provides a prime example of an area where climate change and human activity are rapidly transforming its biogeography. Recent advances in time series analysis, computational resources, and remote sensing data have enabled more comprehensive and nuanced mapping of ecosystem dynamics at regional scales (*40*–*43*). These advances provide the means to reduce the uncertainty in the trajectory of forest dynamics across the state. Here we build upon this past work to develop a multidecadal (1985–2021) time series of vegetation type cover fractions and disturbance type across California. We use these datasets to investigate the following questions: (1) How has the distribution of forest cover shifted across Californian ecoregions over the last 37 years? (2) To what extent are these observed changes driven by fire, harvest, and die-off? (3) How has tree cover change been related to climate? This work will provide the quantification of tree cover dynamics necessary to more effectively administer California’s climate action policies and to better understand how temperate ecosystems are being transformed by climate change.

## Results

### Net changes in vegetation cover across California

Tree cover was highest in northern California along the coast and interior mountain regions at mid and high elevations (Figure 1a). By ecoregion (Supplementary Figure S1), mean tree cover was highest in the Northern Coast and Mountains (41%) and lowest in the Southern Coast and Mountains (15%) (Supplementary Table S1). In contrast, shrub cover was present, but not dominant, throughout all ecoregions and herbaceous cover was dominant in the Central Coast and Foothills ecoregion.

**Figure 1.**
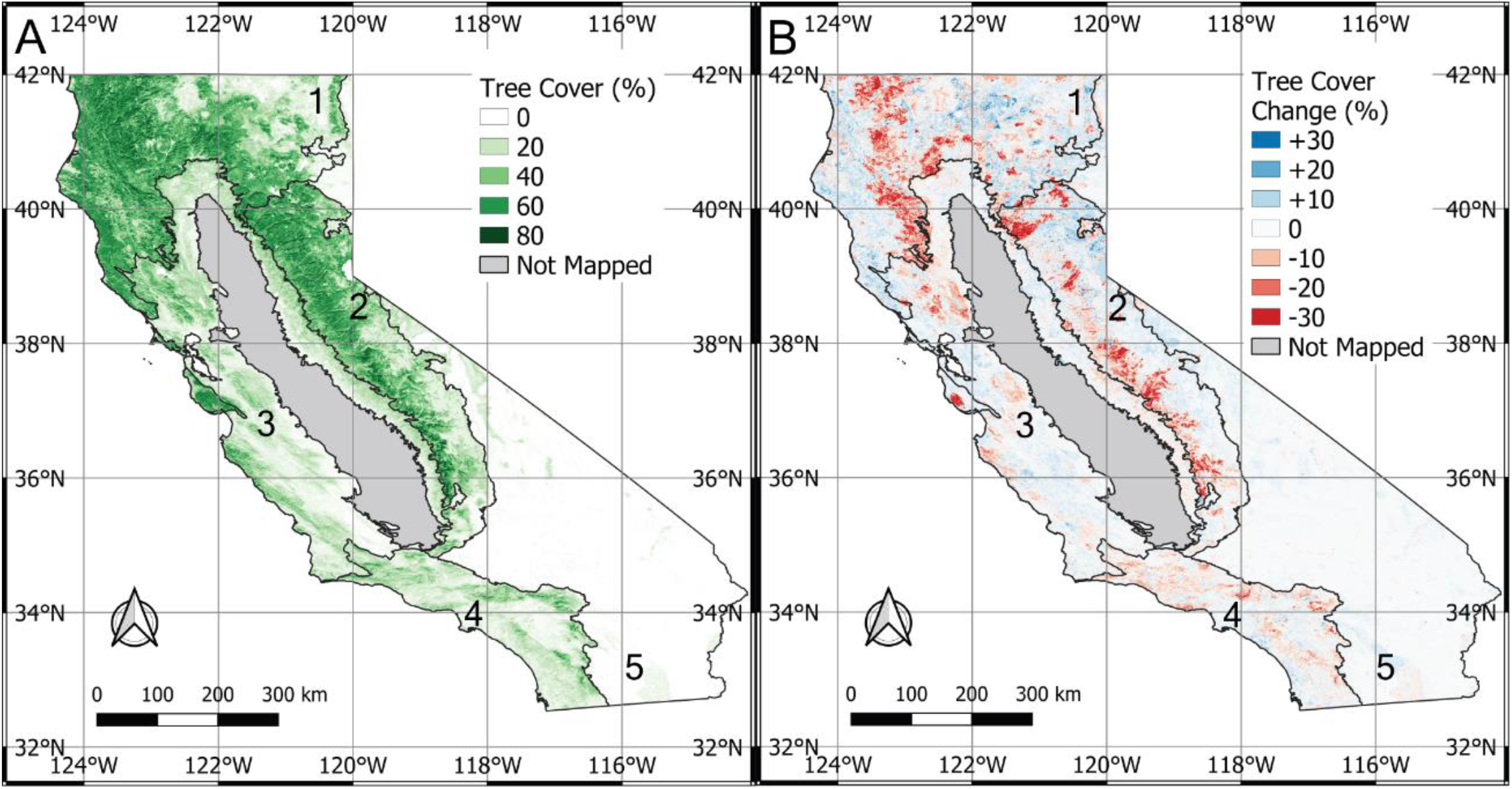
Distribution of tree canopy cover and changes in tree cover across California. a) Tree canopy cover in 1985. b) Change in tree cover between 1985 and 2021. Because we excluded pixels mapped as agricultural land cover, we exclude from our maps the Central Valley ecoregion, since it is composed primarily of agricultural land cover. Lines indicate the boundaries of the aggregated ecoregions, and numbers indicate ecoregions: 1 is the North Coast and Mountains, 2 is the Sierra Nevada, 3 is the Central Coast and Foothills, 4 is the South Coast and Mountains, and 5 is the Deserts. See Supplementary Figure S1 for more information.

From 1985 to 2021, the total tree cover area of California declined by about - 3783 km^2^ or about 5.5% relative to initial cover area (Table 1). In contrast, shrub and herbaceous cover area generally increased between 1984 and 2021, with a net gain of 3371 km^2^ (+6.2%) and 1642 km^2^ (+3.0%), respectively, while bare ground cover decreased by 1278 km^2^ (−3.1%). Note that these changes do not include land cover changes in the Desert or Central Valley ecoregions. As described below, there was considerable variability within individual ecoregions.

**Table 1.** Ecoregion-specific changes in vegetation cover and their drivers from 1985 to 2021. Values listed are in units of km^2^, which represents the cover area estimated using equation 1. Parenthetical values are in units of proportion (%) relative to the initial total cover area within each ecoregion in 1985. The initial totals of vegetation cover area in 1985 are provided in Supplementary Table S1. In areas where multiple disturbances occurred in sequence, the change in tree cover area reported are those corresponding to the disturbance type associated with the most recent disturbance.

| Ecoregion | Δ Tree | Δ Shrub | Δ Herb | Δ Bare | Δ Tree in no disturbance | Δ Tree in fires | Δ Tree in harvest | Δ Tree in die-off |
| --- | --- | --- | --- | --- | --- | --- | --- | --- |
| North Coast and Mountains | -983 (-3.1) | +1044 (+5.7) | +803 (+7.6) | -863 (-6.7) | +900 | -1498 | -320 | -99 |
| Sierra Nevada | -1406 (-7.3) | +517 (+4.5) | +1104 (+19.8) | -267 (-1.7) | +301 | -1384 | -241 | -166 |
| Central Coast and Foothills | -777 (-6.2) | +1,619 (+11.0) | -497 (-1.6) | -345 (-4.4) | +177 | -743 | -18 | -3 |
| South Coast and Mountains | -617 (-11.6) | +191 (+2.0) | +229 (+3.3) | +197 (+3.7) | -57 | -562 | -24 | -3 |
| California | -3783 (-5.5) | +3371 (+6.3) | +1642 (+3.1) | -1278 (-3.1) | +1321 | -4187 | -587 | -246 |

Severe drops in tree cover occurred in localized patches throughout the Northern Coast and Mountains, Sierra Nevada, and Southern Coast and Mountains ecoregions (Figure 1b). Severe forest losses occurred at both fine scales (e.g., patches < 1 km^2^), primarily associated with timber harvest operations, and at coarse scales (e.g., patches > 100 km^2^), primarily associated with recent large wildfires (Figure 2). Fires were distributed throughout all the ecoregions, whereas timber harvests were concentrated in the North Coast and Mountains ecoregion and forest die-off was concentrated in the Sierra Nevada ecoregion (Supplementary Table S1). There was also considerable spatial variability of disturbance within ecoregions, with fires in the North Coast and Mountains ecoregion primarily affecting the Klamath Mountains and fires in the South Coast and Mountains ecoregion primarily affecting inland mountain areas.

**Figure 2.**
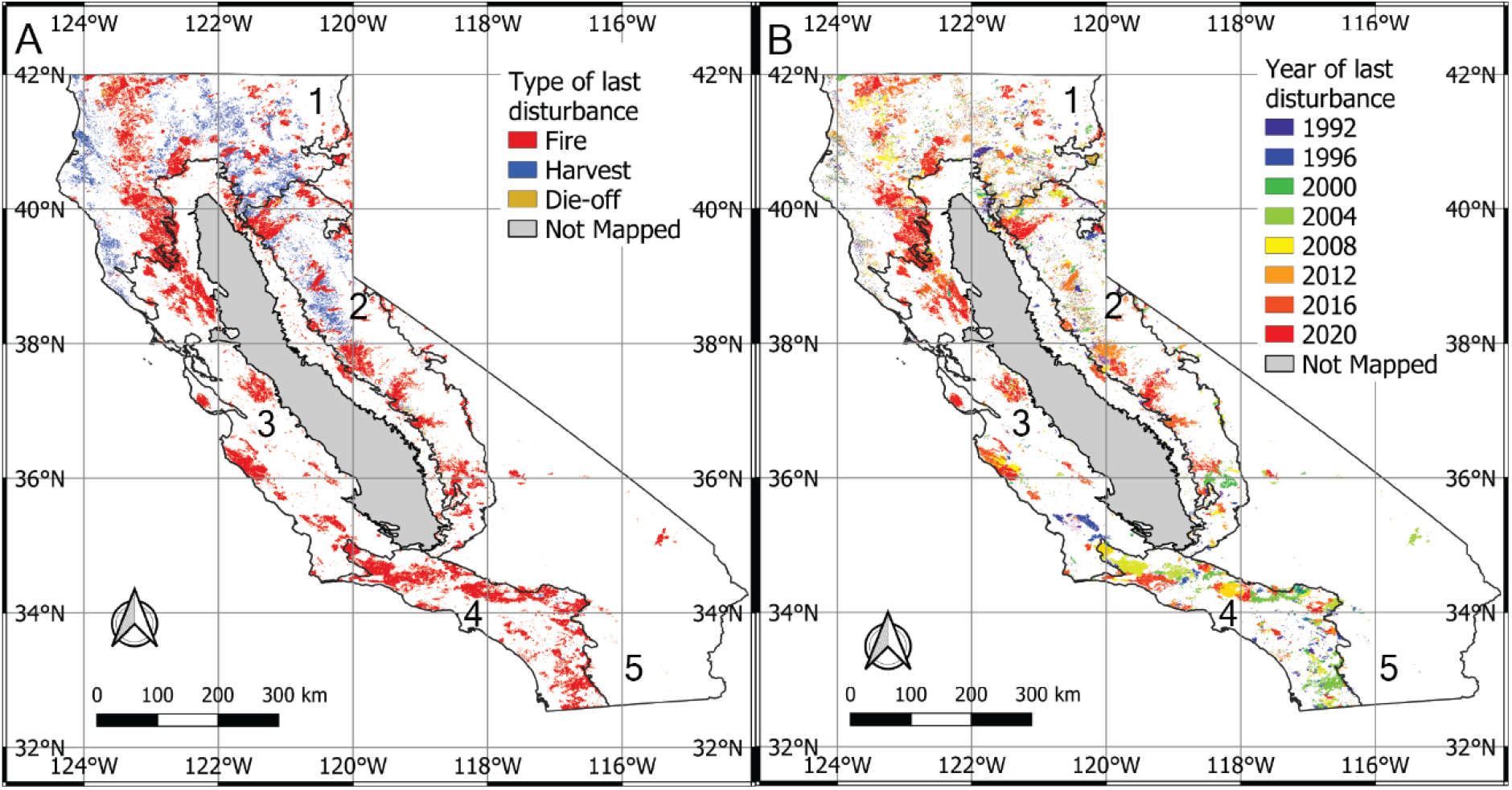
Type and timing of the most recent disturbance throughout California. a) Disturbance type (last disturbance mapped at each location in the record) and b) timing of the last disturbance. Because we excluded pixels mapped as agricultural land cover, we exclude from our maps the Central Valley ecoregion, since it is composed primarily of agricultural land cover. Lines indicate the boundaries of the aggregated ecoregions, and numbers indicate ecoregions: 1 is the North Coast and Mountains, 2 is the Sierra Nevada, 3 is the Central Coast and Foothills, 4 is the South Coast and Mountains, and 5 is the Deserts. See Supplementary Figure S1 for more information.

Loss of tree cover occurred at all latitudes in the state and primarily at middle elevations (i.e., between 700 and 2000 m), except in the southern part of the state (i.e., south of 35°N) where tree cover loss was largely pervasive regardless of elevation (Figure 3). Gains in tree cover occurred primarily north of 35°N at low elevation (i.e., below 1,000 m) and at high elevations (above 2000 m), particularly in the North Coast and Mountains. Tree-cover loss at mid-elevations coincided with high levels of fire, harvest, and die-off (Figure 4). Harvest occurred primarily in the North Coast and Mountain and Sierra Nevada ecoregions at low and middle elevations. Die-off occurred primarily at middle elevations of the North Coast and Mountains and Sierra Nevada ecoregions. Fire was the predominant disturbance in the Central Coast and Foothills and South Coast and Mountains ecoregions.

**Figure 3.**
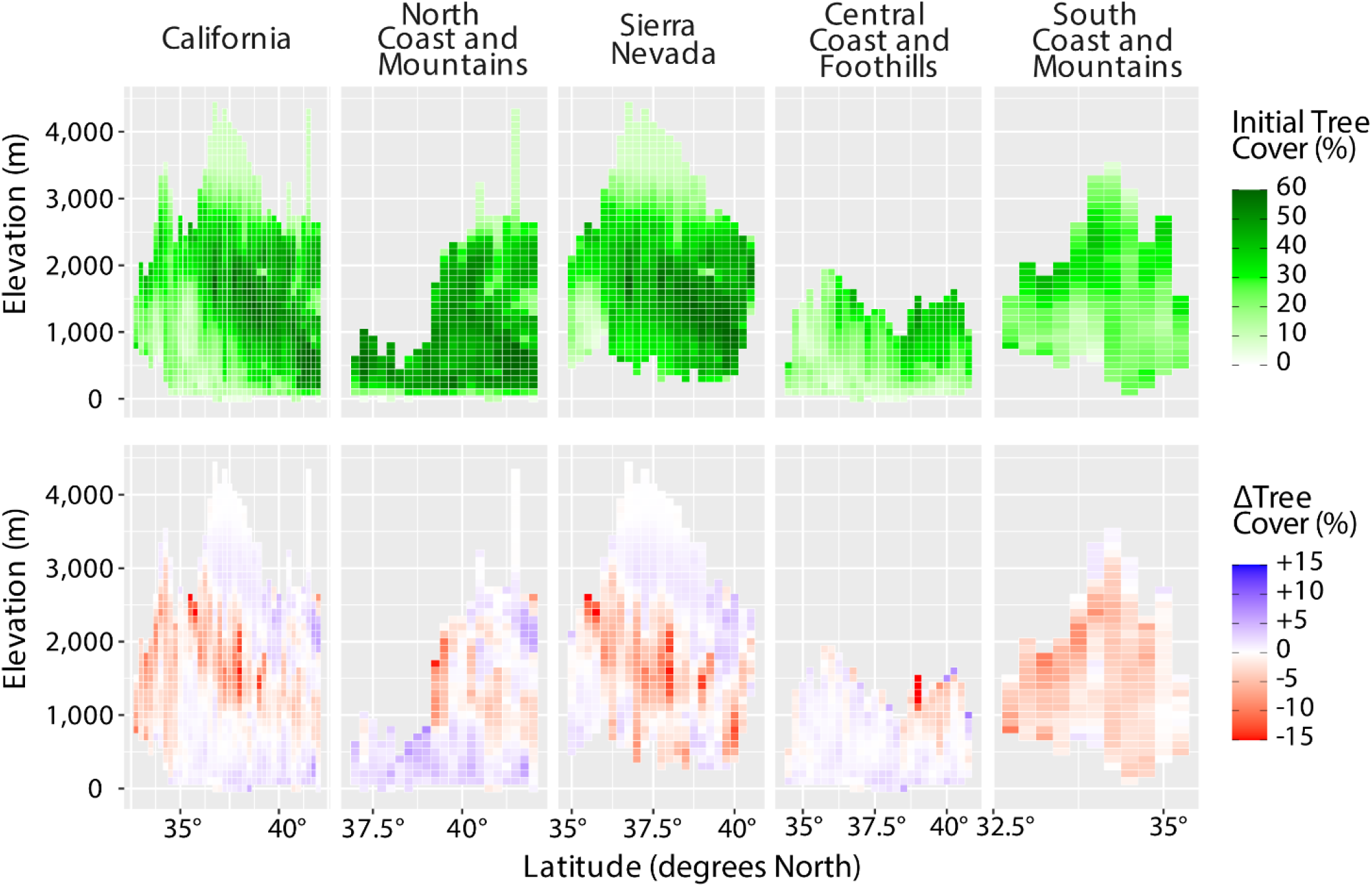
Latitudinal and elevational distribution of California’s tree cover. Top row: Tree cover in 1985. Bottom row: net change (2021 minus 1985) as a function of elevation and latitude. Average tree cover in each elevation-latitude bin was determined by calculating the TCA within each bin and dividing by the land surface area of each bin.

**Figure 4.**
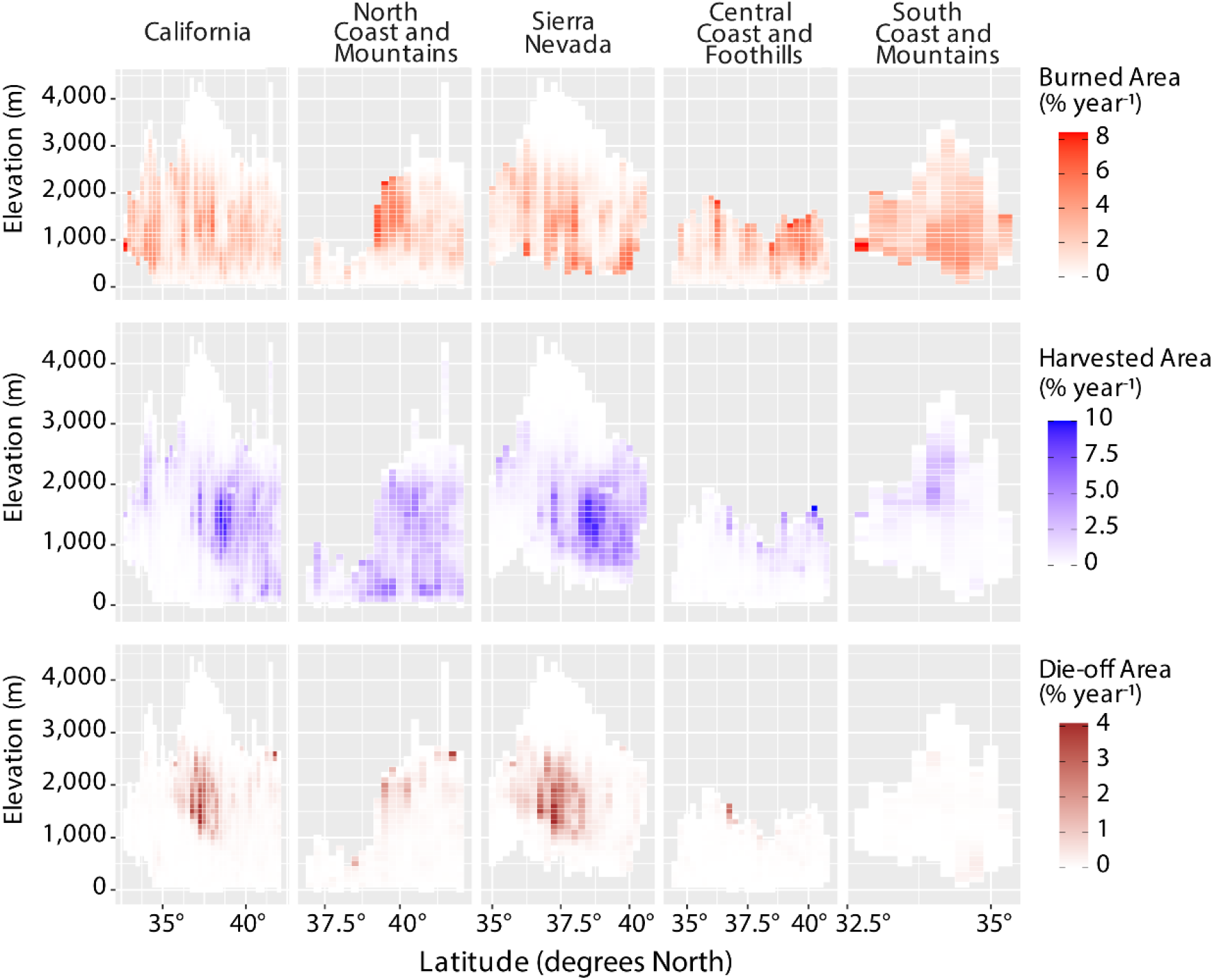
Disturbance as a function of ecoregion, elevation, and latitude averaged during 1985 - 2020. Rates shown are for all areas within each elevation – latitude bin, regardless of vegetation composition. Top row: fire. Middle row: harvest. Bottom row: die-off. Percent changes are estimated by normalizing the total disturbance area by the total land surface area in each bin (and excluding crop, water, and urban pixels).

### Temporal dynamics in vegetation cover

From 1985 to 2021, statewide changes in vegetation cover area varied considerably on annual and decadal timescales. While tree cover area generally decreased over the study period, there was an initial period of tree cover area expansion (+1423 km^2^, or +2.0% of initial tree cover area) during 1985–1999 that was counteracted by an even larger magnitude decline during the last two decades (−5206 km^2^, or -7.5% from 2000–2021) (Figure 5a). Shrub cover area, in contrast, increased nearly monotonically from 1985 to 2019 (+5349 km^2^, or +10.0%) before suddenly declining between 2019 and 2021 (−1978 km^2^, or -3.7%). The sharp declines in tree and shrub cover area between 2019 and 2021 coincided with extensive fires in 2020.Bare ground area had a negative trend, and both bare ground and herbaceous cover area increased sharply at the end of the period as a following large fire years in 2018 and 2020 (Figure 5a).

**Figure 5.**
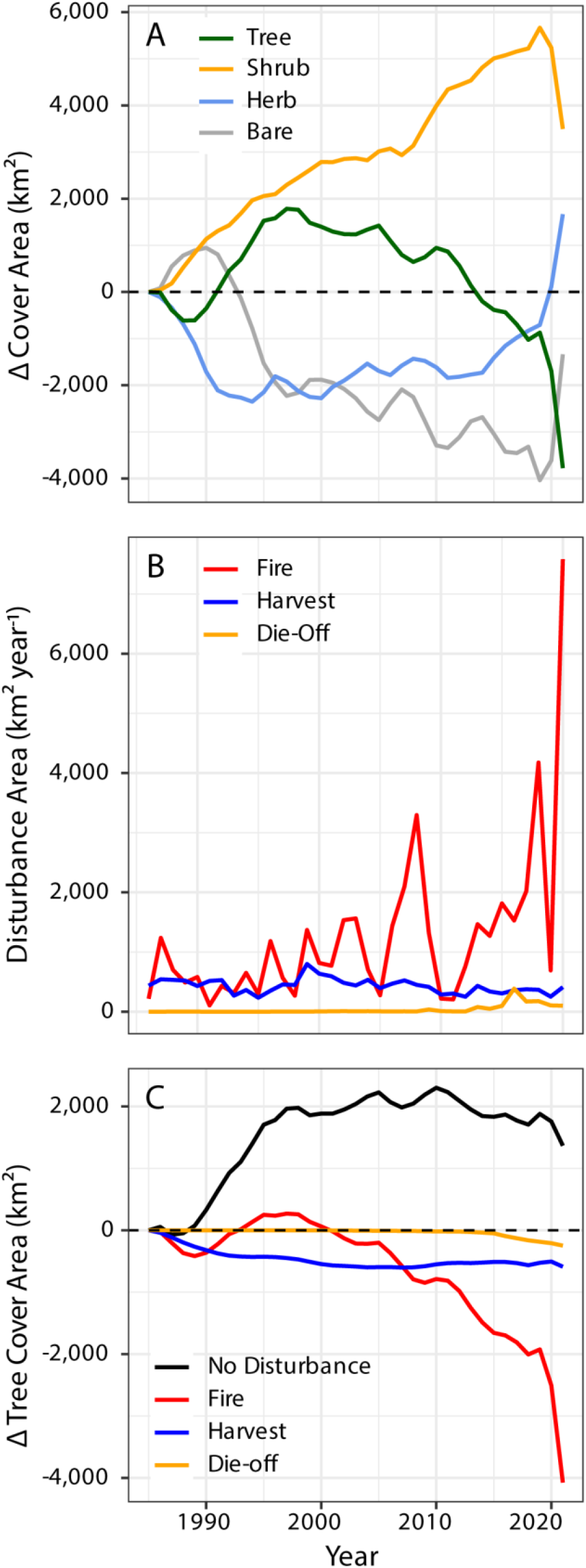
Temporal dynamics of vegetation cover, disturbance, and attribution of tree cover change to disturbance loss and recovery across California. a) Cumulative change in cover area from 1985– 2021. b) Estimates of annual disturbance area for the three mapped disturbance types from 1985 - 2020. Disturbances in 2021 were not mapped because Landsat data are not yet available for the fall of 2021. c) Total change in tree cover area occurring in areas experiencing each of the disturbance types. The individual lines in panel c sum to the dark green line in panel a.

Disturbance was the primary driver of changes in tree cover area during the study period (Figure 5b). Considering the different agents of tree cover loss, fire was the most important driver (4187 km^2^) followed by harvest (587 km^2^) and die-off (246 km^2^) (Table 1). Disturbance thus collectively accounted for a loss of 5020 km^2^ of tree cover area (7.3% of initial tree cover area), which was partially counteracted by 1321 km^2^ of tree cover gains in areas outside of disturbances (1.8% of initial tree cover area) (Figure 5c). Harvest was the dominant driver of tree cover area loss during the first two decades and was replaced by fire during the last two decades (Figure 5c). When interpreting the tree cover area time series (Figure 5), it is important to remember that changes in tree cover area may originate from changes in spatial extent of forest area as well as changes in leaf area and stand density as a consequence of climate and disturbance drivers.

The interval of tree cover area expansion early in the time series from 1985-1999 coincided with low levels of disturbance and above average levels of annual precipitation. Burned area was lower during 1986–1998 (541 ± 95 km^2^ year^-1^; 95% confidence interval indicates temporal variability) than the long-term mean (1210 ± 462 km^2^ year^-1^). Similarly, the period 1993 - 1998 had a reduced average harvest rate of 353 ± 39 km^2^ year^-1^, which was lower than the average harvest rates for the preceding six-year period 1986 – 1993 of 512 ± 16 km^2^ year^-1^ or for the entire 1985-2020 period of 429 ± 19 km^2^ year^-1^. At the same time, precipitation was about 31% higher (1169 ± 146 mm year^-1^) than the mean during the full study period (889 ± 48 mm year^-1^) (Supplementary Figure S2). Above average precipitation likely increased tree cover area as a consequence two different mechanisms. First, it is consistent with anomalous low levels of burned area, since higher levels of precipitation are likely to increase live fuel moisture, and thus reduce fire occurrence and rates of fire spread (*44*). Second, sustained periods with above average precipitation generally increase tree growth and leaf area, especially in dry forests such as those in California and may enhance rates of tree seedling recruitment in areas with a mixture of vegetation types (*45*).

Tree cover declined nearly monotonically after the year 2000, owing in part to the occurrence of very large fire years in 2008, 2018, and 2020, when fires accounted for 3294 km^2^, 4175 km^2^, and 7583 km^2^ of burned area, respectively. At the same time, the drought during 2012–2016 increased the incidence of drought-induced tree mortality, contributing to a loss of tree cover area of 385 km^2^ year^-1^ in 2016. The incidence of wildfire and drought in the 2000s and 2010s resulted in large decreases in tree cover in mountainous areas. In contrast, harvest rates had a decreasing trend during the latter two decades of the time series in ecoregions where harvest was significant (North Coast and Mountains and Sierra Nevada), suggesting that the state-wide decline in tree cover area was not primarily harvest-driven.

### Geographic variability in vegetation dynamics

Every ecoregion experienced a net loss in tree cover, particularly as a result of the large 2020 fire year (Figure 6a). Though all ecoregions lost tree cover to fire, by far the largest proportional change in tree cover occurred in the South Coast and Mountains ecoregion, which experienced a 1.5% cumulative loss in average tree cover area across the ecoregion. Compared to initial levels of average tree cover in 1985 (14.7%), this is equivalent to a 10.1% relative loss. Most ecoregions experienced modest gains in tree cover during the early part of the study period, but increasing fire more than offset these gains during the decades of the 2000s and 2010s, burning large parts of each ecoregion area (up to 4% in a single year), killing many trees, and shifting more landscape areas into an early successional state where shrubs are dominant (Figure 6b). Following large fire years, shrub and herbaceous cover area increased considerably, though some of the gains in shrub cover in the Central Coast and Foothills ecoregion occurred prior to the large fire years that were observed in the latter part of the record.

**Figure 6.**
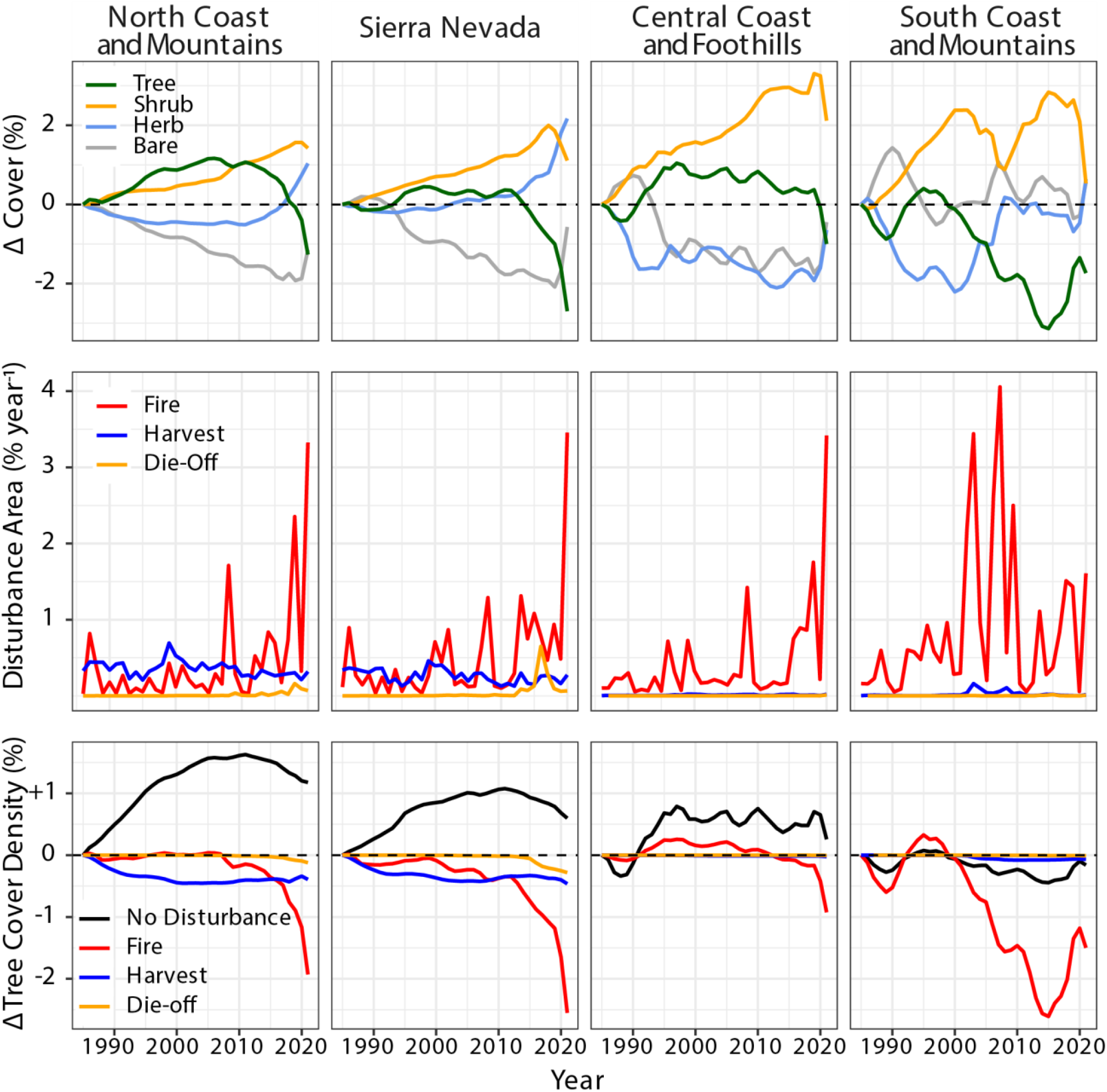
Temporal dynamics of vegetation cover, disturbance, and attribution of tree cover change to disturbance loss and recovery across each ecoregion. To account for different sized areas of each ecoregion, cover fractions are shown as the average across the entire ecoregion area. Top row: Cumulative average vegetation cover change for the four mapped vegetation types from 1985–2021, estimated by dividing cover area for each cover type (equation 1) by the total land surface area in that ecoregion. Middle row: Estimates of annual disturbance for the three mapped disturbance types from 1985 - 2020. Bottom row: Cumulative change in average tree cover occurring in areas experiencing each of the disturbance types. In each column, the lines in panel c sum to the dark green line in panel a.

In contrast, harvest occurred over a steady, but relatively modest, area (0.25– 0.5%), mostly in the North Coast and Mountains and the northern part of the Sierra Nevada ecoregions. In both of these ecoregions, harvest area decreased significantly from 1999–2020, with a trend of -9.1 ± 2.4 km^2^ year^-1^ (p <0.001) for the North Coast and Mountains and -4.1 ± 2.4 km^2^ year^-1^ (p <0.004) for the Sierra Nevada. Forest die-off also occurred primarily in the North Coast and Mountains and Sierra Nevada ecoregions, with substantial episodes of forest die-off occurring primarily in the later part of the time period (from 2012–2018 in the Sierra Nevada and starting in 2017 in the North Coast and Mountains) that coincide with the occurrence of a severe drought (*17, 33*). While forest die-off only occurred in limited areas and timespans, it affected a similar area as fire across the Sierra Nevada during 2016 (Figure 6b).

Outside of areas that experienced disturbance (Figure 6c), all ecoregions except the South Coast and Mountains had increasing levels of tree cover. Expansion of tree cover area is expected in undisturbed areas as a consequence of recovery from fire and harvest prior to the beginning of the Landsat record. The lack of any visible recovery in the South Coast and Mountains ecoregion may reflect some combination of 1) low rates of disturbance in the decades prior to the onset of the Landsat era, reducing the area of post-disturbance recovery, 2) the effects of increasing stress that limits tree establishment and growth in previously disturbed areas, and 3) little or no expansion of tree cover in new areas.

### Sensitivity of tree cover and disturbance to climate

Mean tree cover was highest in areas where mean summer temperature was between 12.5°C and 22.5°C and where mean annual precipitation exceeded 1000 mm year^-1^ (Figure 7a and b). The large majority of tree cover area, which also accounts for available land surface area in each climate bin, occurred in areas with mean summer temperatures between 15 and 25°C and in areas with mean annual precipitation between 500 and 1500 mm year^-1^ (Figure 7c and d). Some of these parts of the climate space exhibit somewhat low tree cover, suggesting that areas with relatively low tree cover make up a relatively large proportion of the state’s total tree cover area.

**Figure 7.**
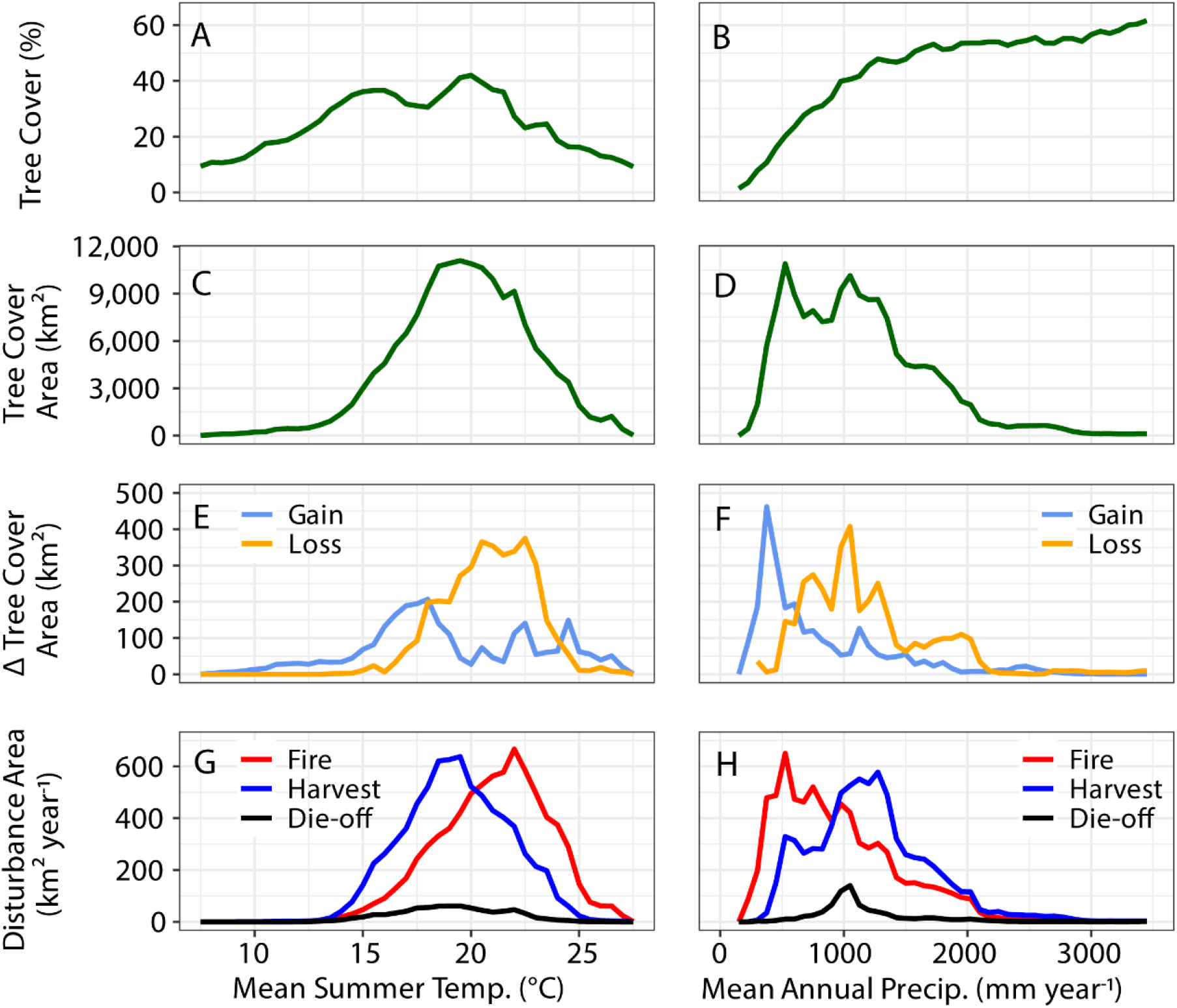
Distribution of initial tree canopy cover and net tree cover change as a function of long-term climate. First column: mean summer temperature. Second column: long-term mean annual precipitation. In second and third rows, areas in km^2^ indicate accumulated changes in tree cover area. a) Variability in initial tree cover as a function of mean summer temperature. b) Variability in initial tree cover as a function of mean annual precipitation. c) Variability in initial tree cover area as a function of mean summer temperature. d) Variability in initial tree cover area as a function of mean annual precipitation. e) Occurrence of tree cover area gains and losses as a function of mean summer temperature. f) Occurrence of tree cover gains and losses as a function of mean annual precipitation. g) Occurrence of fire, harvest, and die-off as a function of mean summer temperature. h) Occurrence of fire, harvest, and die-off as a function of mean annual precipitation. For initial tree cover fraction and area distributions we report our estimates for 1985, the first year of our time series.

Losses of tree cover area in California occurred disproportionately in warmer regions. Tree cover area losses peaked in areas where summer temperatures were between 19 and 23°C (Figure 7e), which nearly coincided with the peak in burned area at 22.5°C (Figure 7g). In contrast, the peak in tree cover area gains occurred at lower summer temperatures between 15 and 18°C, in a climate zone where fire was less prevalent. As a function of the rate of warming, tree cover losses occurred in areas that had faster rates of warming (> 0.04°C year^-1^), which also had high rates of fire. In contrast, net tree cover gains occurred in more coastal regions that had slower rates of warming, or were cooling (Figure 8, Supplementary Figure S3).

**Figure 8.**
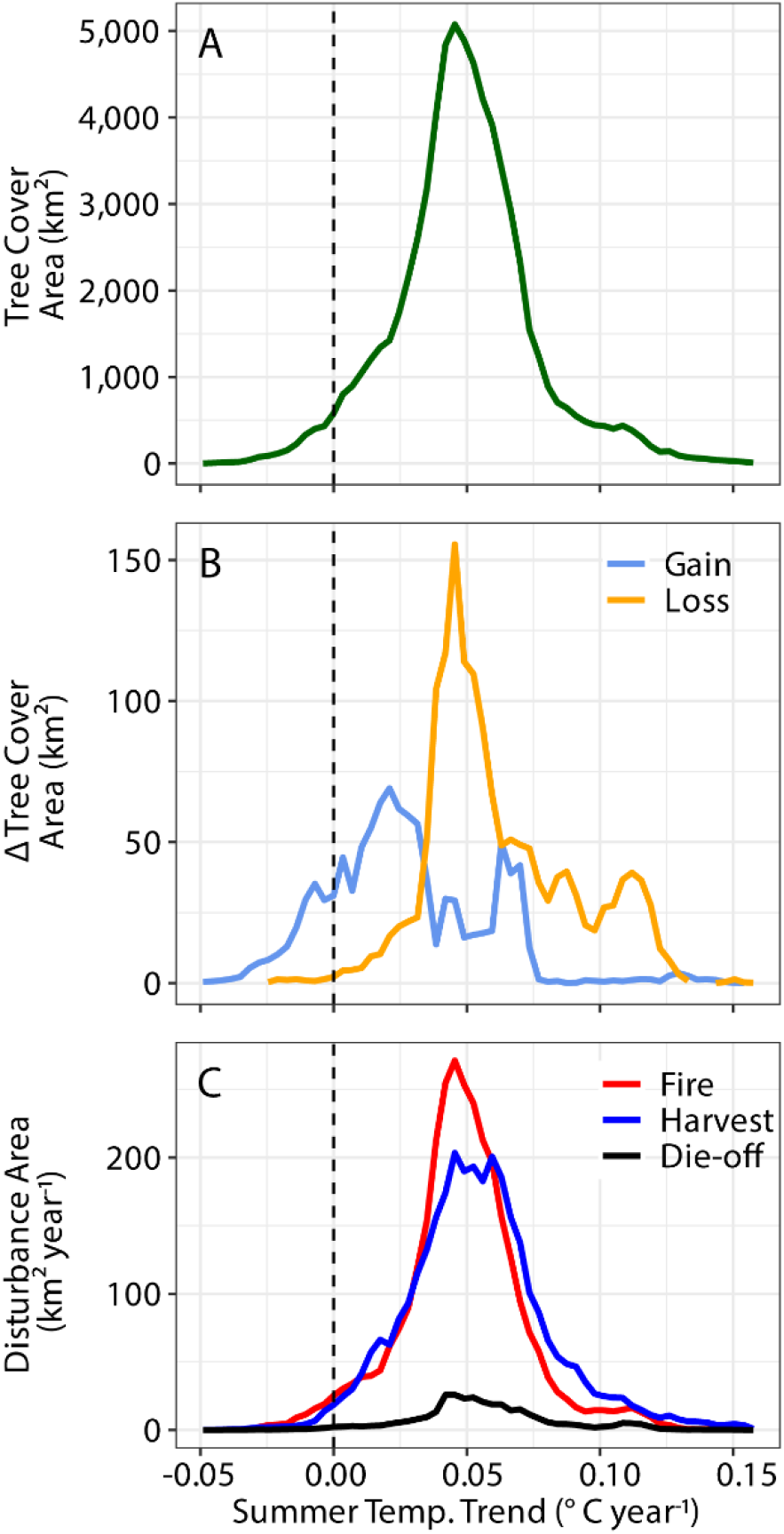
Distribution of tree cover and disturbance dynamics as a function of summer temperature trends across all of California. In panels a and b, areas in km^2^ indicate accumulated changes in both areal extent and sub-pixel density in tree cover. The vertical dashed line indicates regions of 0 trend in summer temperature. See Supplementary Figure S3 for information on the spatial distribution of these trends. a) Variability in initial tree cover density (1985). b) Occurrence of net tree cover gains and losses. c) Occurrence of fire, harvest, and die-off.

As a function of precipitation, tree cover area losses were highest between 700 and 1500 mm year^-1^ whereas tree cover gains were highest in drier areas with less than 700 mm year^-1^ (Figure 7f). Tree cover area losses occurred in areas where initial tree cover is quite high, suggesting a high level of vulnerability in dense forests. The gains in tree cover area occurred in places where initial tree cover was quite low, with values less than half of the levels supported in moister climates (Figure 7b). Burned area was also concentrated in drier regions, with a peak around 500 mm year^-1^ and overlapping considerably with areas that had both significant losses and gains in tree cover area.

Drought-induced die-off of forests occurred most prominently in a narrow band of precipitation centered around 1000 mm year^-1^ primarily in the Sierra Nevada. One hypothesis for the structure of the gains and losses as a function of precipitation is that conifer forests, which thrive at higher levels of precipitation, were disproportionately contributing to tree cover losses, whereas the gains are occurring as a consequence of the expansion of hardwood species in drier areas. This hypothesis is consistent with past work by McIntyre *et al.* (2015), who showed that, during the 20^th^ century, areas in California experiencing relatively high moisture deficits were transitioning from more pine-dominated ecosystems to more oak hardwood dominance. Because we found that the vast majority of precipitation trends were not significant (at p = 0.05 level, see Supplementary Figure S3), we ultimately did not analyze changes in tree cover relative to trends in precipitation.

## Discussion

### Tree cover loss and implications for natural climate solutions

We used time series of Landsat imagery and machine learning to show tree cover has substantially declined across California over the 37-year period from 1985 through 2021, primarily as a consequence of increasing fires and drought during the last decade. Tree cover area declined by 5.5% for the state as a whole, with the most severe losses occurring in the south and in middle elevation mountain regions. The overall spatial pattern suggests a broad-scale, fire-driven redistribution, shifting tree cover area from interior mountains to foothills and coastal areas. Global environmental change may be driving enhanced growth in undisturbed areas through mechanisms such as CO_2_ fertilization (*46*). However, warming generally tends to lead to either drought-induced mortality or increased fire risk (*19*) and experimental evidence suggests that the accumulation of net primary production in tree biomass may not be enhanced by increased CO_2_ concentrations (*47*).The net loss of tree cover documented here and underlying datasets we used to assess these changes have several significant implications for the use of forests as natural climate solutions.

First, our study provides quantitative estimates of the impact of increasing fire activity on forest status. Fires were the main driver of tree cover losses. Given expected future fire risks are likely to intensify (*48, 49*), our work suggests that an increasing share of California’s dense and productive forests will experience significant fire-driven declines. California’s carbon offsets program provides carbon credits to landowners who manage their forest resources for improved carbon sequestration. These programs include a “buffer pool”, a set of carbon credits that are set aside to account for the potential of reversals, the intentional or unintentional release of carbon back to the atmosphere due to storms, fire, and other factors (*50*). Our work suggests that the buffer pool set aside to account for fire losses may need to be reassessed in order to effectively account for the increasing risk that fires and other disturbances pose to temperate forests.

Second, the densest forests appear particularly vulnerable to changing disturbance and high temperatures (Figure 7), and the risks to these forests appear to be increasing due to warming (Figure 8). To be effective, natural climate solutions should focus on reducing future losses of already present carbon in these areas, rather than efforts to enhance carbon storage. Warming, vulnerable areas may not be suitable or efficient areas for improved forest management projects, including mid-elevation forest in the Sierra Nevada and forests in the mountains of Southern California. This is especially important as climate change is projected to reduce biomass densities in these areas (*51*). In contrast, the forests in the northern coast of California appear to have expanding tree cover where the effects of climate-driven changes in wildfires are not yet having an effect. These are also the densest forests in California. Wildfire protection and careful management of these northern forests may yield more effective carbon mitigation potential.

Third, both the tree cover fraction and disturbance products developed here may be helpful for monitoring of the effectiveness of offset projects. Development and deployment of these remote sensing-based monitoring systems will provide unbiased, independent verification of changes in harvest rates or forest growth relative to business-as-usual scenarios. An important next step with respect to assessment of forest offset projects is to develop an aboveground biomass carbon product using Landsat and lidar remote sensing data that is internally consistent with the tree cover and disturbance products described here. This product would allow the quantification of carbon stocks lost in wildfires or sequestered by improved forest management, providing a means to either optimize placement of carbon offset projects or to verify the change in carbon in existing projects. These products would also allow researchers to estimate how climate change is influencing rate of regrowth to better understand California’s total carbon budget and how fire protection programs translate to saved carbon and improved post-fire recovery.

### Disturbance and climate change drive tree cover dynamics

The incidence of disturbance and climate change is reshaping the composition of ecosystems across California. Fires drove the large majority of tree cover loss, and climate warming has led to an increase in burned area in California (*39*). The loss of trees promoted an increase in shrub cover as a consequence of post-disturbance recovery and succession. In many areas, shrubs that establish early in succession are likely to be replaced by trees, regenerating the forests that were originally lost. However, with the increasing trend in fire disturbance observed here, post-disturbance recovery appears unable to keep pace and California may be facing lasting biome shifts (*52*). Simultaneously, increasing disturbance causes the average forest stand age to decline, which may result in significant changes in ecosystem function (*53*). Climate change also may be modifying successional trajectories and the ability of trees to reestablish in disturbed areas (*54*). In the South Coast and Mountains ecoregion, the lack of evidence for tree cover expansion in areas outside of known disturbance provides indirect evidence for a possible change in succession and reduced efficiency of forest recovery.

We observed a considerable increase in tree cover area in the early part of the record in northern ecoregions, which coincided with a period where rates of disturbance were lower and the climate was wetter than average. It is likely that this increase in tree cover occurred by means of densification of existing forested areas in response to increased precipitation and as a consequence of lower disturbance rates from fire and harvest. Compared to the relatively high rates of forest harvest in the 1950s and 1960s (*55*), relatively low rates of forest harvest in the 1980s and 1990s may have led to an increase in tree cover. Changes in forest harvest practices in the 1990s may also have been driven by a change in forest policy related to the conservation of the spotted owl (*56, 57*), suggesting that changes in human-induced forest management can have significant, measurable impacts on the availability of forest resources. The high rates of harvest in the decades prior to the satellite record would result in otherwise unattributed tree cover gains as harvested forests recovered. In addition, the increases in tree cover may reflect the occurrence of structural overshoot, whereby rapid growth in response to temporarily more favorable conditions results in a buildup of tree biomass exceeding the long-term average that typical climatic environment may support (*25*). The increases in leaf area and stand density can result in increased vulnerability to drought, resulting in later self-thinning, or a buildup of fuels that results in more severe fires (*58*). Indeed, the increased tree cover area from the early part of the study is more than compensated for by severe fires and drought that occur later in the time series, which may be a delayed response to the extra growth in the 1990s. It is unclear to what degree structural overshoot as opposed to warming-induced changes in disturbance regimes are driving these temporal dynamics, but it is clear from the temporal structure that there is a recent shift in the environmental factors controlling California’s ecosystems.

Losses of tree cover were most pronounced in warmer parts of the state, whereas cooler parts of the state experienced gains in tree cover. Notably, net tree cover losses were much more prominent in areas experiencing summer warming greater than 0.3^°^C year^-1^ (Figure 8). This suggests that climate change is contributing to a large-scale restructuring of tree cover that is likely to continue with future warming. These losses occurred in areas of significant warming even beyond the areas that experienced substantial fires or harvest, suggesting that undetected drought-induced die-off and other climate stressors are driving significant tree cover loss in the state. As California’s summers have been warming rapidly (Supplementary Figure S2), the climate space that these ecosystems occupy will also shift, potentially increasing the risk of fire, die-off, or other climate-induced stress to a wider proportion of the state’s trees.

### Implications for California’s total carbon budget

A key policy goal for California is the management of its greenhouse gas budget, as stipulated in the California Global Warming Solutions Act of 2006. The state is committed to improving carbon sequestration in the state’s land via, in part, the management of its forest resources (*7*). However, considerable uncertainty remains in the quantification of the carbon budget, with differing methodologies yielding divergent results. Estimates based on combining field plots measurements with remote sensing data show large losses of carbon from California’s forests (−5.3 ± 1.6 Tg C year^-1^) (*6*), while estimates relying solely forest inventory data without spatially explicit information suggest large gains of carbon (+6.5 ± 0.6 Tg C year^-1^) (*7*). Our results suggest that if carbon is accumulating in aboveground live biomass at a statewide-level, it must be occurring at a high enough rate in the coastal areas of the north that the significant tree cover losses we observe in the southern part of the Sierra Nevada ecoregion, the interior parts of the North Coast and Mountains Ecoregion, and throughout the South Coast and Mountains ecoregion are more than offset. The development of a multi-decadal carbon product, when coupled with the tree cover and disturbance products described here, has the potential to reconcile these differences.

### Key uncertainties and future directions

In this study, we established a remote sensing method for detecting multiple types of disturbance in a common framework. However, as with many remote sensing-based products, it can be challenging to detect and attribute disturbances that are less severe. Our machine learning algorithm was trained primarily on the most intense forest management and drought-induced mortality events, but there are many instances of forest management practices and mortality events that only induce minor changes in tree cover. For example, forest management practices that focus on understory removal, or uneven-aged silviculture treatments of the forest, will alter the coverage of tree canopy, but may not be detected by our algorithm. Our indication of the contribution of disturbance to tree canopy change is therefore a lower bound, as we generally do not capture the potentially widespread low-intensity changes. This is particularly relevant for drought-induced mortality and low intensity management - while we capture the most severe mortality events, our maps miss many of the less intense mortality events. This discrepancy may explain potentially the relatively low accuracy of drought-induced die-off detection. Our mapping is focused on the most intense areas and weighted by forest cover loss intensity, and more work is necessary to refine the inputs and sensitivities of such a disturbance detection algorithm to identify drought-induced die-off in a timely manner.

### Conclusions

California is at the forefront of forest management policies to counteract rapid climate change, preserve carbon stocks, improve water quality, and mitigate wildfire risk. Understanding changes in vegetation cover is critical for targeting ecosystems for efficient management and for predicting where fuels build-up and wildfire risk is high, as well as quantifying the impact of various management strategies. In this study, we have found that California’s ecosystems are becoming increasingly heterogeneous in age and composition due to increasing disturbance, and that disturbances pose a significant threat to the integrity of forests in all ecosystems across the state. The warmest parts of the state are losing substantial tree cover and experiencing limited recovery, while many cooler areas are experiencing considerable increases in tree – climate warming will exacerbate these vulnerabilities. Net gains and losses of tree cover area were associated with spatial variability in climate, and increased climate warming threatens the integrity of forests. The drivers and consequences of these vegetation dynamics need to be explored in detail in order to prioritize forest conservation and management resources across the state. The results will improve understanding the recent history of California’s forests and quantification of California’s carbon budget, and ultimately provide insights into the vulnerability of natural climate solutions in many fire-prone and rapidly warming ecosystems. Better attribution of increases in vegetation cover will be necessary to craft more efficient and equitable forest carbon offset programs.

## Materials and Methods

We combined existing geospatial datasets of disturbance events and vegetation cover with Landsat-derived metrics using machine learning, effectively extrapolating the original snapshot datasets to create spatiotemporally continuous datasets over the entire state of California. Analyses and datasets are subdivided into ecoregions based on aggregations of the Environmental Protection Agency level 3 Ecoregions (Supplementary Figure S1; Table 1; Supplementary Table S2) (*59*). Vegetation cover and disturbance were mapped for all ecoregions except for the Central Valley, which is composed primarily of agricultural land. For the subsequent analyses, we exclude the Central Valley and Desert ecoregions given that most of these lands had minimal tree cover. Instances where “California” is used as an ecoregion represents the sum total of the state of California, but still excluding the Central Valley and Desert ecoregions as indicated in Supplementary Figure S1.

### Change detection in remote sensing time series

We used time series of surface reflectance data from the Landsat collection of Earth observation satellites, including Landsat 4 and 5 Thematic Mapper, Landsat 7 Enhanced Thematic Mapper+, and Landsat 8 Operational Land Imager for 1985 - 2021 (*60*). Specifically, we used the U.S. Geological Survey’s Collection 1 Surface Reflectance product and quality-screened for clouds, sensor saturation, and other data quality issues. To estimate the timing and magnitude of disturbance, we temporally segmented the time series of each spectral band at each 30 m pixel using the Continuous Change Detection and Classification (CCDC) algorithm (*42*). The temporal segmentation fits piecewise harmonic regression models to identify the timing and location of statistically significant breaks. The algorithm generates a database of model coefficients for producing smoothed, interpolated ‘synthetic’ reflectance data at each pixel. Model breaks and synthetic reflectance were inputs to the machine learning workflow used to map plant cover and disturbance. Temporal segmentation was performed using Google Earth Engine (*61*).

### Modeling fractional vegetation cover

To model fractional vegetation cover, we trained a machine learning algorithm on existing datasets of vegetation from the Multi-Resolution Land Characteristics (MRLC) Consortium, including the National Land Cover Database (NLCD) and the Rangeland Condition Monitoring and Projection (RCMAP) datasets. The NLCD provides maps of land cover type and fractional cover of trees (woody plants taller than 5 m) across the CONUS approximately every half decade from the year 2001 (*62–64*). We used the “analytical” version of the tree cover dataset and simplified the dominant land cover dataset classification scheme (Supplementary Table S3). The RCMAP product maps fractional cover of various shrub (woody plants less than 5 m), herbaceous (grasses, forbs, and cacti), and bare cover (exposed soil, sand, and rocks) components for just the rangelands in the western US for the years 1985-2018 (*65*). RCMAP additionally maps shrub sub-types, litter, and non-photosynthetic vegetation. The MRLC’s fractional cover data are originally generated with a machine learning approach using Landsat spectral metrics as inputs and trained on manually labelled high-resolution imagery. Note that these datasets are mapped over a limited temporal and spatial domain. Specifically, the NLCD land cover and tree cover datasets are snapshots of the years 2001, 2006, 2011, and 2016, while RCMAP data only cover the rangelands, rather than the entirety of California. To fully analyze the spatial and temporal variability of different vegetation types across California, it was necessary to develop a machine learning approach for extrapolating these data through time. To simplify our maps, we do not map litter, and our model thus predicts only live vegetation cover or bare ground.

To develop our fractional plant cover time series dataset, we used synthetic reflectance from the CCDC models as inputs to four random forest regression models (*66, 67*), one each to predict fractional cover of tree, shrub, herbaceous, or bare ground. We additionally trained a random forest classification model to predict land cover type in order to filter our dataset for water, agriculture, and urban areas. To develop the model, we drew a simple random sample of 1.5M pixels across the NLCD and RCMAP 2016 datasets, which had relatively higher accuracy than the earlier datasets (*68*). We held out 20% (0.3M pixels) of this dataset for testing, and trained each model on the remaining 1.2M pixels. At each sampled pixel, we used the CCDC outputs to model the surface reflectance of 5 spectral bands, as well as 7 spectral vegetation and tasseled cap indices (Supplementary Table S4). The inputs for fractional cover mapping included spectral values modeled for both mid-summer (August 1, Julian day of year 213) and spring (March 1, Julian day of year 60) intervals for the year 2016. Because the CCDC-derived models effectively interpolate values of surface reflectance between observations, we are able to consistently use specific dates for our reflectance values rather than identifying temporal metrics (e.g. maximum) within a seasonal range. We used spring and summer values because California’s ecosystems respond to a Mediterranean climate, which has mild and wet springs when shrubs and herbaceous leaf out and hot, dry summers when trees remain leafed but most shrubs and herbaceous vegetation senesce. These seasons capture the major phenological variability between the vegetation types in California, allowing us to partition the drought deciduous herbaceous vegetation from the evergreen trees and shrubs. Additionally, we extracted the elevation from the US Geological Survey ⅓ arc-second National Elevation Dataset at each sampled pixel. The resulting random forest regression models were used to predict and map the annual fractional cover of tree, shrub, herbaceous, and barren land across California.

We modified the resulting vegetation cover predictions with several post-processing steps. The RCMAP data from Rigge et al. (2021) exhibited considerable bias with their independent validation samples. To rectify this, we calibrated our predictions of shrub, herbaceous, and bare cover using the linear regressions reported by Rigge et al.(2021), improving the dynamic range of the predictions. Additionally, since individual RF models predicting fractional cover were independent of one another, the sum of fractional cover predicted did not always sum to 100% (average vegetation cover sum was 97 ± 8.9% across all pixels, mean and one standard deviation). As total cover, by definition, must equal 100%, we normalized each pixel’s fractional plant cover fractions such that they collectively added to 100%. To do this, we divided the prediction of each vegetation type by the sum of all the predicted cover fractions within a pixel for each year. We applied each of the five random forest models to predict time series of land cover type and fractional plant cover for each year in 1985 - 2021 across all areas of California. For this study, we do not analyze changes in vegetation cover in the Central Valley or the Deserts (Supplementary Figure S1). We used land cover predictions to mask areas that were ever water, agriculture, and urban areas from 1984-2021 from our analyses. Using these models, we produce maps of the tree, shrub, herbaceous, and barren cover density at each 30 m pixel in California for each year from 1984 – 2021. Note that early successional trees that grow after fire or harvest are likely to be classified as shrubs in our algorithm when their height is less than 5 m.

### Characterizing vegetation cover area

To characterize regional sums of vegetation cover, we report sums of total vegetation cover in km^2^ throughout this study for individual ecoregions and for the state as a whole. We do this by using the following equation:

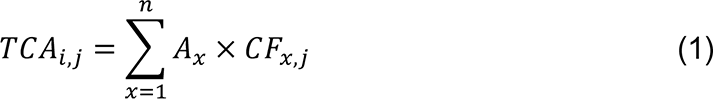

where TCA_i,j_ is the total vegetation cover area for ecoregion i and vegetation cover type j, A_x_ is the surface area of each 30 m pixel x (900 m^2^), n is the total number of pixels within each ecoregion, and CF_x,j_ is the cover fraction (unitless) at each pixel x for cover type j, up to the nth pixel in each region. To provide ecoregion averages for each cover fraction, we divide TCA_i,j_ by the total land surface area within each ecoregion (excluding urban, agricultural, and water areas). In this study, we refer to this TCA value as the vegetation cover area (e.g. tree cover area or shrub cover area). We refer to the fractional cover as the vegetation cover (e.g. tree cover).

We note that changes in tree cover fraction at each pixel may originate from the expansion or contraction of leaf area of existing trees within the pixel as well as changes in the number of trees (woody vegetation taller than 5 m) within the pixel. Periods of favorable or less favorable climate may modify tree cover fraction by either mechanism, whereas fire and harvest primarily modify tree cover by means of the latter mechanism. At an ecoregion level, tree cover area may change from the lateral expansion or contraction of areas that have trees, or by a change in the leaf area or stand density of areas that have trees to begin with. Our definition of tree cover fraction is consistent with previous work creating continuous vegetation cover fraction products from AVHRR and MODIS satellite imagery (*69–71*) and has the advantage of being able to detect more subtle changes in tree cover caused by partial mortality events such as those associated with drought or insects. Our results may differ significantly from studies that only account for categorical classifications of forest versus non-forest land at each 30m pixel (e.g. using categorical classifications from Landfire) (*72*), since the forest lands we observe generally have tree cover fractions lower than 100% and our ecoregion sums are represent the sum of tree cover for all the pixels within the domain, including sub-pixel changes.

### Modeling and mapping disturbance agents

To better understand potential drivers of changes in vegetation cover described above, we generated a unified, consistent time series map of ecosystem disturbance agents across California. We mapped the timing and location of these disturbances by classifying CCDC model breaks according to disturbance causal agents. CCDC model breaks were classified as belonging to one of five classes - fire, harvest, die-off, greening, or browning. Fire, harvest, and die-off are considered primary disturbances of interest. Greening and browning are not considered disturbances, but are indicative of subtler changes in ecosystem condition (i.e. related to climate variability or post-disturbance regrowth) that trigger a statistical break in the CCDC temporal segmentation. We trained the classifier using a reference dataset of disturbances identified from the intersection of CCDC model breaks and existing geospatial datasets of disturbance.

We developed the reference dataset of disturbance agents by sampling CCDC model breaks (n = 150000) from across California and the 1985-2020 time period and classifying them as belonging to one of five classes on the basis of their spatial and temporal overlap with existing geospatial archival databases. We did not estimate disturbance information for 2021 because the data for the complete year was not yet available (in contrast, vegetation cover requires data up to August, which was covered when this study was conducted, and so vegetation cover for 2021 was mapped). We held out 20% (n = 25000) of the model breaks as a test dataset for evaluating the model. For geospatial information on fires, we referred to the California Department of Forestry and Fire Protection (CAL FIRE) Fire and Resource Assessment Program (FRAP) dataset, which provides the most complete set of fire perimeters in California since 1900 (*73*). For information on timber harvests, we used data from CAL FIRE’s Timber Harvest Plans (*74*) (THP), which provides perimeters for permitted timber harvest projects. For drought-induced die-off, we used the US Forest Service’s Insect and Disease Detection Survey (*75*) (IDS), which provides annual polygons describing approximate areas experiencing insect and drought-induced die-off as observed from aircraft. Because remote sensing approaches can struggle to detect canopy-scale changes that are of low intensity or that primarily affect the understory, we restricted the THP and IDS polygons used to those experiencing the most intense disturbance. For THP data, we used polygons having intensity that was medium or high (*56*), and for IDS data, we used polygons having mortality falling into the most severe class, where 30-50% of trees are observed to have died. CCDC model breaks that fell within the perimeter of these disturbances and had a break date within one year of the archival dataset’s reported disturbance were labeled as being caused by that disturbance. CCDC model breaks that did not occur within these perimeters were considered land changes that were not associated with disturbance and were classified as greening or browning depending on the sign of the break’s change magnitude (e.g., a positive change in NDVI was considered greening, and a negative change was considered browning).

Using the disturbance reference dataset, we trained the random forest model to attribute disturbance type using the change magnitude of each spectral band at each CCDC model break. The change magnitude depends on the size of the residuals in between the pre-break model and the post-break observations and is calculated for each spectral band. The change magnitudes identified substantial variability in the spectral signature of the different disturbance types and provides the basis for model training and inference (Supplementary Figure S4). The disturbance model was then applied across the study domain to predict the timing and type of disturbance, including areas beyond the original dataset perimeters. Additionally, we used elevation and slope as inputs. The minimum mapping unit was 8100 m^2^.

### Performance of vegetation and disturbance models

The main goal of our machine learning approach was to extrapolate the fractional cover databases for the year 2016 across all of California over the timespan 1985 - 2021. The regression models predicted fractional plant cover with high skill, with an R^2^ between modeled and reference values ranging from 0.80 to 0.95 and RMSE ranging from 4.4 to 9.6 % cover using the held-out test dataset (Supplementary Figure S5). Tree cover mapping had an R^2^ of 0.85 and an RMSE of 4.4 % cover, shrubs had an R^2^ of 0.80 and an RMSE of 9.6% cover, herbaceous had an R^2^ of 0.9 and an RMSE of 9.4% cover, and bare ground had an R^2^ of 0.95 and an RMSE of 8.4 % cover. There was generally very little bias in the models across all the plant cover types, as estimated as the slope of the line fit between the modeled and reference data, except for a slight underestimation of tree cover at high values (Supplementary Figure S5).

The classification model for disturbance type also had relatively high accuracy, with an overall accuracy of 84% when compared to a 20% test dataset (n = 25000). However, there was considerable variability in the accuracy of mapping different disturbance types. For predicting fire, the user’s and producer’s accuracy were relatively high (82% and 89%), respectively (Supplementary Table S5). The fire maps tended to estimate lower areas of burning across California relative to estimates from FRAP, estimating 56 ± 17% lower rates of burn per year. This is largely because FRAP assumes complete burning within fire perimeters, but our maps included sub-perimeter heterogeneity in burns, allowing for unburned islands and spatial detail that is a common feature of fires in temperate ecosystems (*26, 27, 36*). Additionally, our map captured 7716 km^2^ of additional fire beyond FRAP perimeters. In contrast to fire, harvest mapping had somewhat lower accuracy, with a user’s accuracy of 72% and a producer’s accuracy of 81% (Supplementary Table S5). Part of the challenge with accurately predicting forest activity is the uncertainty in the archival records. Knight et al. inspected time series of aerial photos of timber harvest polygons in the CAL FIRE database. They detected modest spatial and temporal biases in this data. Finally, mapping of drought-induced die-off was comparatively low, with a producer’s accuracy of 50% and a user’s accuracy of 78% (Supplementary Table S5). This may lead to an underestimation of the area impacted by drought-induced die-off.

### Relationships between vegetation and climate

We analyzed the distribution of tree cover, changes in tree cover, and disturbance frequency as a function of long-term mean climate. We used precipitation and mean temperature data from the PRISM Climate Group for 1984 through 2020 (*76*). We averaged mean tree cover, net changes in tree cover area, and incidence of disturbances according to summer temperature and annual precipitation bins (0.5^°^C and 50 mm). Annual precipitation was summed according to the water year. We similarly analyzed the distributions of these properties as a function of elevation (100 m bins) and latitude (0.25^°^ bins).

To analyze the sensitivity of tree cover change to climate change, we also analyzed trends in mean summer temperature and annual precipitation. We estimated these trends at each pixel using linear regression, with year as the predictor and mean summer (June, July, August) temperature or total annual precipitation (for the water year) as the response. To interpret trends in vegetation cover relative to climate change, we binned tree cover and disturbance data into summer temperature trend and annual precipitation trend bins (0.01°C year^-1^ and 10 mm year^-1^, respectively).

## Acknowledgments

We gratefully acknowledge grant support from California’s Strategic Growth Council (SGC) Climate Change Research Program to M. Goulden at the Center for Ecosystem Climate Solutions and from the University of California’s National Laboratories (UCNL) Laboratory Fees grant program to J. Randerson and M. Goulden (as a part of the California Ecosystems Futures project). Additional support was provided by UC Irvine to the Geospatial Data Solutions Center. Randerson acknowledges additional long-term support from NASA, including grants from Carbon Monitoring System and Modeling Analysis and Prediction Programs, and from the U.S. Dept. of Energy Office of Science Biological and Environmental Research RUBSICO Science Focus Area. The authors declare that they have no competing interests. All data needed to evaluate the conclusions in the paper are present in the paper or the Supplementary Materials. The disturbance and vegetation maps can be provided by J.A.W. upon request. J.A.W. and J.T.R. designed the study. J.A.W., C.K., M.L.G., and J.B.B. generated the disturbance data set. J.A.W. generated the vegetation cover dataset and conducted data analysis. J.A.W. and J.T.R. wrote the first draft of the manuscript. All authors contributed to the interpretation of the results and editing of subsequent manuscript drafts.

## Supplementary Tables

**Supplementary Table S1.** Description of the area, climate, land cover, and disturbance for each ecoregion in our study. Vegetation areas indicated represent the initial total areas for the state in 1985, and parentheses indicate the total of each ecoregion covered by that vegetation type. Areas do not add to 100% because of the exclusion of water land cover (e.g. lakes and rivers). For fire, harvest, and mortality areas, error ranges indicate 95% confidence intervals over the temporal variability. Our study does not include the Central Valley ecoregion, and the California ecoregion also excludes that area.

| Ecoregion | Area (km<sup>2</sup>) | Precip. (mm year<sup>-1</sup>) | Temp (°C) | Tree Area (km<sup>2</sup>) | Shrub Area (km<sup>2</sup>) | Herb Area (km<sup>2</sup>) | Bare Area (km<sup>2</sup>) | Fire (km<sup>2</sup> year<sup>-1</sup>) | Harvest (km<sup>2</sup> year<sup>-1</sup>) | Die-off (km<sup>2</sup> year<sup>-1</sup>) |
| --- | --- | --- | --- | --- | --- | --- | --- | --- | --- | --- |
| North Coast and Mountains | 77644 | 1269 ± 114 | 18.4 ± 0.2 | 31658 (41%) | 18250 (24%) | 10563 (14%) | 12782 (16%) | 339 ± 179 (0.4%) | 265 ± 30 (0.3%) | 10 ± 8 (<<0.1%) |
| Sierra Nevada | 52120 | 872 ± 88 | 18.1 ± 0.3 | 19139 (37%) | 11254 (22%) | 5569 (11%) | 15057 (29%) | 246 ± 108 (0.5%) | 130 ± 18 (0.2%) | 21 ± 21 (<<0.1%) |
| Central Coast and Foothills | 76679 | 707 ± 74 | 22.8 ± 0.2 | 12518 (16%) | 14675 (19%) | 30580 (39%) | 7829 (10%) | 326 ± 161 (0.4%) | 10 ± 2 (<<0.1%) | 1 ± 0 (0%) |
| South Coast and Mountains | 35917 | 445 ± 65 | 21.0 ± 0.2 | 5312 (15%) | 9609 (27%) | 6913 (19%) | 5315 (15%) | 299 ± 117 (0.8%) | 8 ± 4 (<<0.1%) | 1 ± 0 (<<0.1%) |
| California | 242360 | 883 ± 94 | 19 ± 0.2 | 68628 (28%) | 53788 (22%) | 53625 (22%) | 40983 (17%) | 1210 ± 462 (0.5%) | 413 ± 47 (0.2%) | 32 ± 26 (<<0.1%) |

**Supplementary Table S2.** Aggregated ecoregions used for this study and the EPA level 3 ecoregions that comprise each. See Supplementary Figure S1 for a map of the aggregated ecoregions.

| Agg. Eco. Code | Aggregated Ecoregion Name | US EPA Level 3 Ecoregion | US EPA Level 3 Code |
| --- | --- | --- | --- |
| **1** | North Coast and Mountains | Coast Range | 1 |
|  |  | Cascades | 4 |
|  |  | Klamath Mountains/California High North Coast Range | 78 |
|  |  | Eastern Cascades Slopes and Foothills | 9 |
| **2** | Sierra Nevada | Sierra Nevada | 5 |
| **3** | Central Coast and Foothills | Central California Foothills and Coastal Mountains | 6 |
| **4** | South Coast and Mountains | Southern California Mountains | 8 |
|  |  | Southern California/Northern Baja Coast | 85 |
| **5** | Deserts | Central Basin and Range | 13 |
|  |  | Mojave Basin and Range | 14 |
|  |  | Northern Basin and Range | 80 |
|  |  | Sonoran Basin and Range | 81 |
| **6** | Central Valley | Central Valley | 7 |

**Supplementary Table S3.** Description of the land cover types, from the National Land Cover Database 2016 (NLCD). The numbers in brackets indicates the data code for each class in the original NLCD dataset.

| **Land Cover** | **NLCD Classes Contained** | **Descriptions** |
| --- | --- | --- |
| Water/Ice | Water [11]; Ice [12] | Areas of open water, generally with less than 25% cover of vegetation or soil. |
| Urban | Developed, Medium Intensity [23]; Developed, High Intensity [24] | Areas with a mixture of constructed materials and vegetation. Impervious surfaces account for at least 50% of the total cover |
| Barren | Barren Land [31] | Areas of bedrock, desert pavement, sand dunes, gravel pits and other accumulations of earthen material. Generally, vegetation accounts for less than 15% of total cover |
| Forest | Deciduous Forest [41]; Evergreen Forest [42]; Mixed Forest [43] | Areas dominated by trees generally greater than 5 meters tall, and greater than 20% of total vegetation cover. |
| Shrub | Shrub/Scrub [52] | Areas dominated by shrubs less than 5 meters tall with shrub canopy typically greater than 20% of total vegetation and tree canopy less than 20% total cover. This class includes true shrubs, young trees in an early successional stage or trees stunted from environmental conditions. |
| Grass | Grassland/Herbaceous [72] | Areas dominated by graminoid or herbaceous vegetation, generally greater than 80% of total vegetation. These areas are not subject to intensive management such as tilling, but can be used for grazing. |
| Crops | Pasture/Hay [81]; Cultivated Crops [82] | Areas of grasses planted for livestock grazing the production of seed or hay crops, or annual crops and woody orchards. Pasture/hay/crop vegetation accounts for greater than 20% of total vegetation. |

**Supplementary Table S4.** Spectral bands and indices from Landsat used for modeling of land cover and disturbance type. Land cover was modeled using the springtime and summertime values for each year, while disturbance type was modeled using the change in summertime values from the year before and the year after a given CCDC break.

| Band or Index | Formula | Reference |
| --- | --- | --- |
| Green | Band 2 (B2) |  |
| Red | Band 3 (B3) |  |
| Near Infrared | Band 4 (B4) |  |
| Short-wave Infrared 1 | Band 5 (B5) |  |
| Short-wave Infrared 2 | Band 7 (B7) |  |
| Normalized Burn Ratio | $NBR = \frac{(B4 - B7)}{(B4 + B7)}$ | (77) |
| Enhanced Vegetation Index | $EVI = 2.5 \cdot \frac{(B4 - B3)}{(B4 + 6 \cdot B4 - 7.5 \cdot B2 + 1)}$ | (78) |
| Normalized Difference Vegetation Index | $NDVI = \frac{(B4 - B3)}{(B4 + B3)}$ | (79) |
| Normalized Difference Moisture Index | $NDMI = \frac{(B4 - B5)}{(B4 + B5)}$ | (17) |
| Tasseled Cap Brightness | $TCB = 0.303 \cdot B1 + 0.279 \cdot B2 + 0.474 \cdot B3 + 0.558 \cdot B4 + 0.508 \cdot B5 + 0.186 \cdot B7$ | (80) |
| Tasseled Cap Greenness | $TCG = -0.289 \cdot B1 - 0.243 \cdot B2 - 0.544 \cdot B3 + 0.724 \cdot B4 + 0.084 \cdot B5 - 0.18 \cdot B7$ | (80) |
| Tasseled Cap Wetness | $TCW = 0.150 \cdot B1 + 0.197 \cdot B2 + 0.328 \cdot B3 + 0.340 \cdot B4 - 0.711 \cdot B5 - 0.457 \cdot B7$ | (80) |

**Supplementary Table S5.**
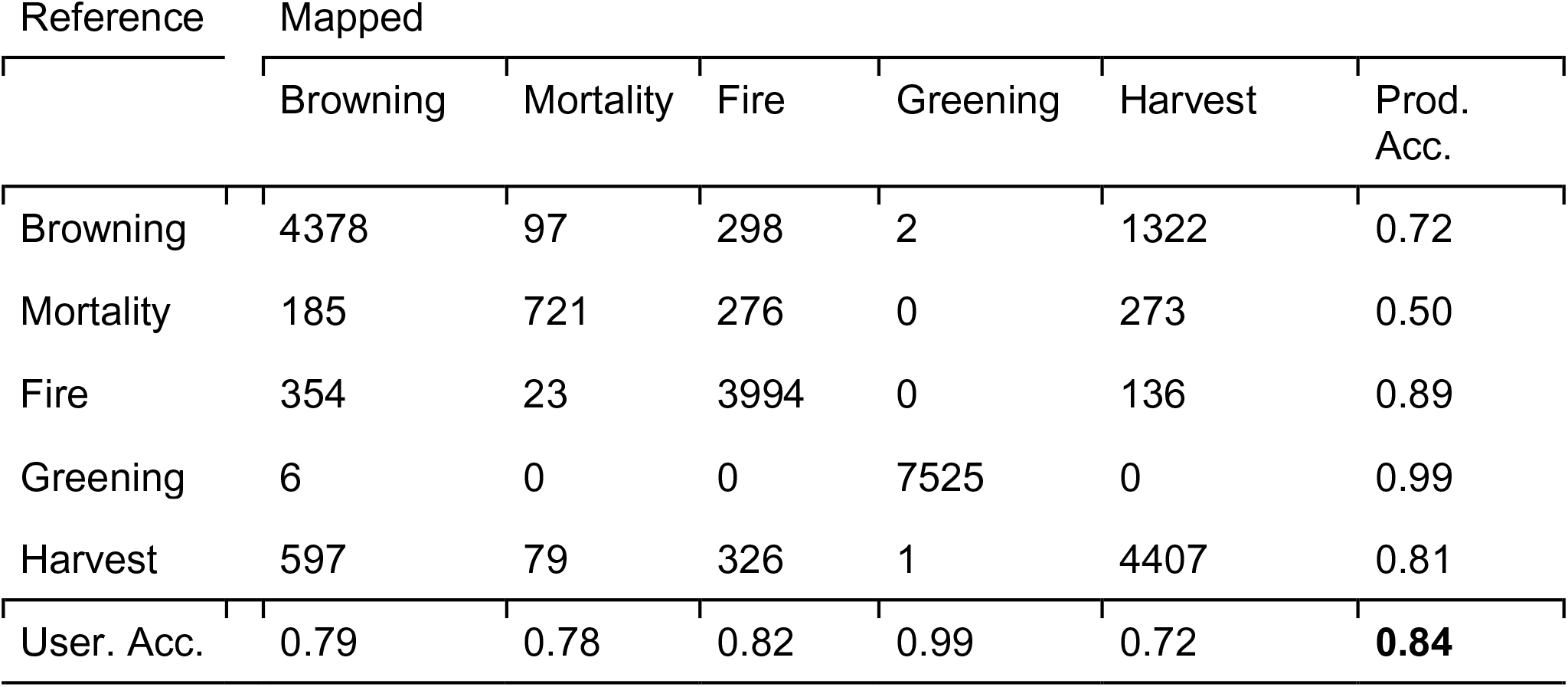
Confusion matrix of the disturbance mapping. Accuracy estimated against a simple random sample across the state of 25000 points. Browning and greening refer to changes in ecosystem state that trigger a model break in CCDC, but are not classified as a disturbance equivalent to the others. Bolded figure indicates the overall accuracy.

| Reference | Mapped |  |  |  |  |  |
| --- | --- | --- | --- | --- | --- | --- |
|  | Browning | Mortality | Fire | Greening | Harvest | Prod. Acc. |
| Browning | 4378 | 97 | 298 | 2 | 1322 | 0.72 |
| Mortality | 185 | 721 | 276 | 0 | 273 | 0.50 |
| Fire | 354 | 23 | 3994 | 0 | 136 | 0.89 |
| Greening | 6 | 0 | 0 | 7525 | 0 | 0.99 |
| Harvest | 597 | 79 | 326 | 1 | 4407 | 0.81 |
| User. Acc. | 0.79 | 0.78 | 0.82 | 0.99 | 0.72 | **0.84** |

## Supplementary Figures

**Supplementary Figure S1.**
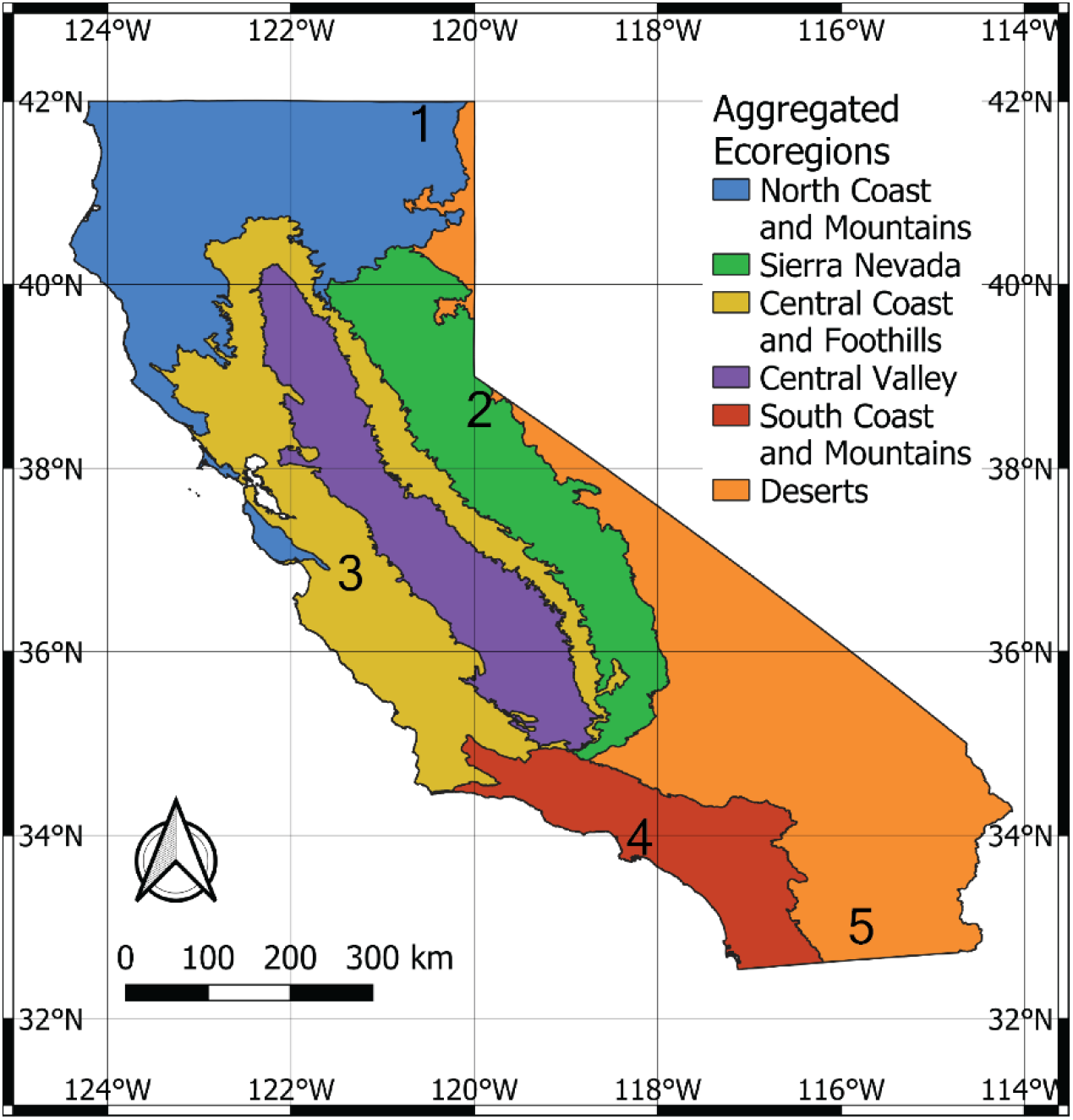
Ecoregions considered in this study. Lines indicate the boundaries of the aggregated ecoregions, and numbers indicate ecoregions: 1 is the North Coast and Mountains, 2 is the Sierra Nevada, 3 is the Central Coast and Foothills, 4 is the South Coast and Mountains, and 5 is the Deserts. As it is not mapped in this study, the Central Valley is not included in any of this study’s maps. See Supplementary Table 1 for details on aggregation.

**Supplementary Figure S2.**
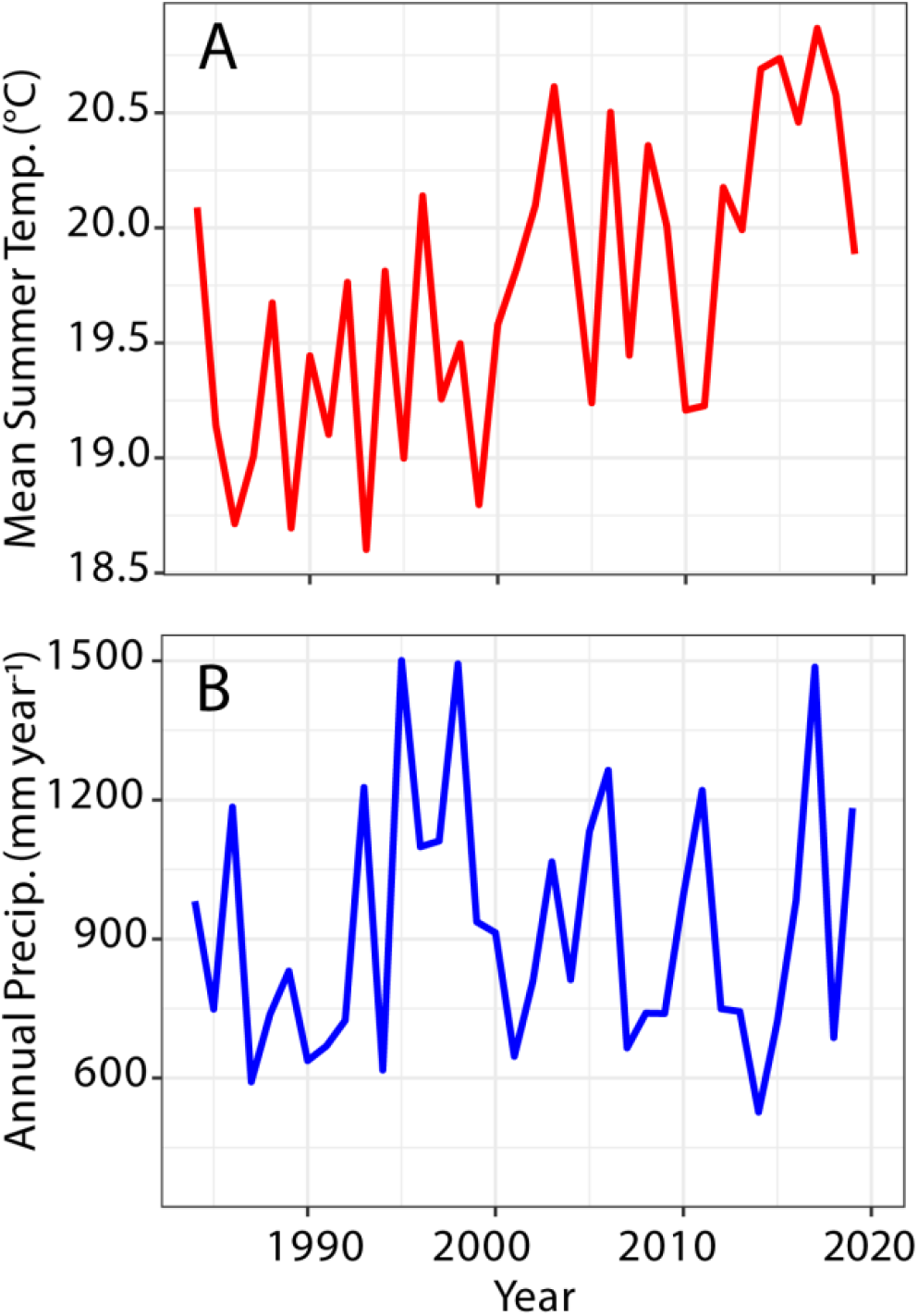
Interannual variability in climate across California, calculated from PRISM and excluding the Central Valley. a) Mean summer temperatures. b) Annual total precipitation. Precipitation represents the water year (from October – September).

**Supplementary Figure S3.**
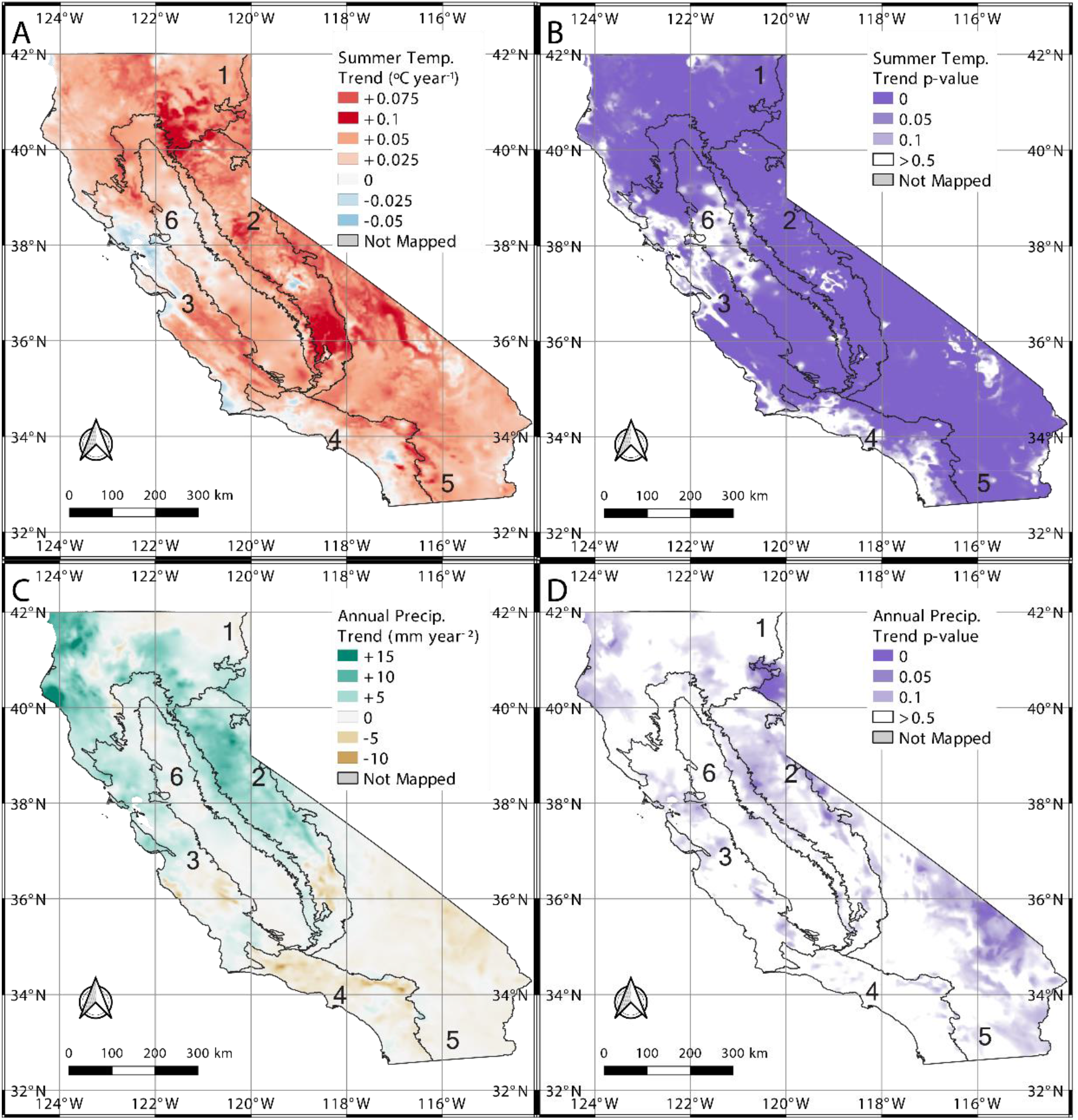
Trends in climate across California, estimated from linear regressions. Lines indicate the boundaries of the aggregated ecoregions, and numbers indicate ecoregions: 1 is the North Coast and Mountains, 2 is the Sierra Nevada, 3 is the Central Coast and Foothills, 4 is the South Coast and Mountains, 5 is the Deserts, and 6 is the Central Valley. a) Trends in summer temperature b) p-value for trends in summer temperature c) Trends in annual precipitation and d) p-value for trends in annual precipitation.

**Supplementary Figure S4.**
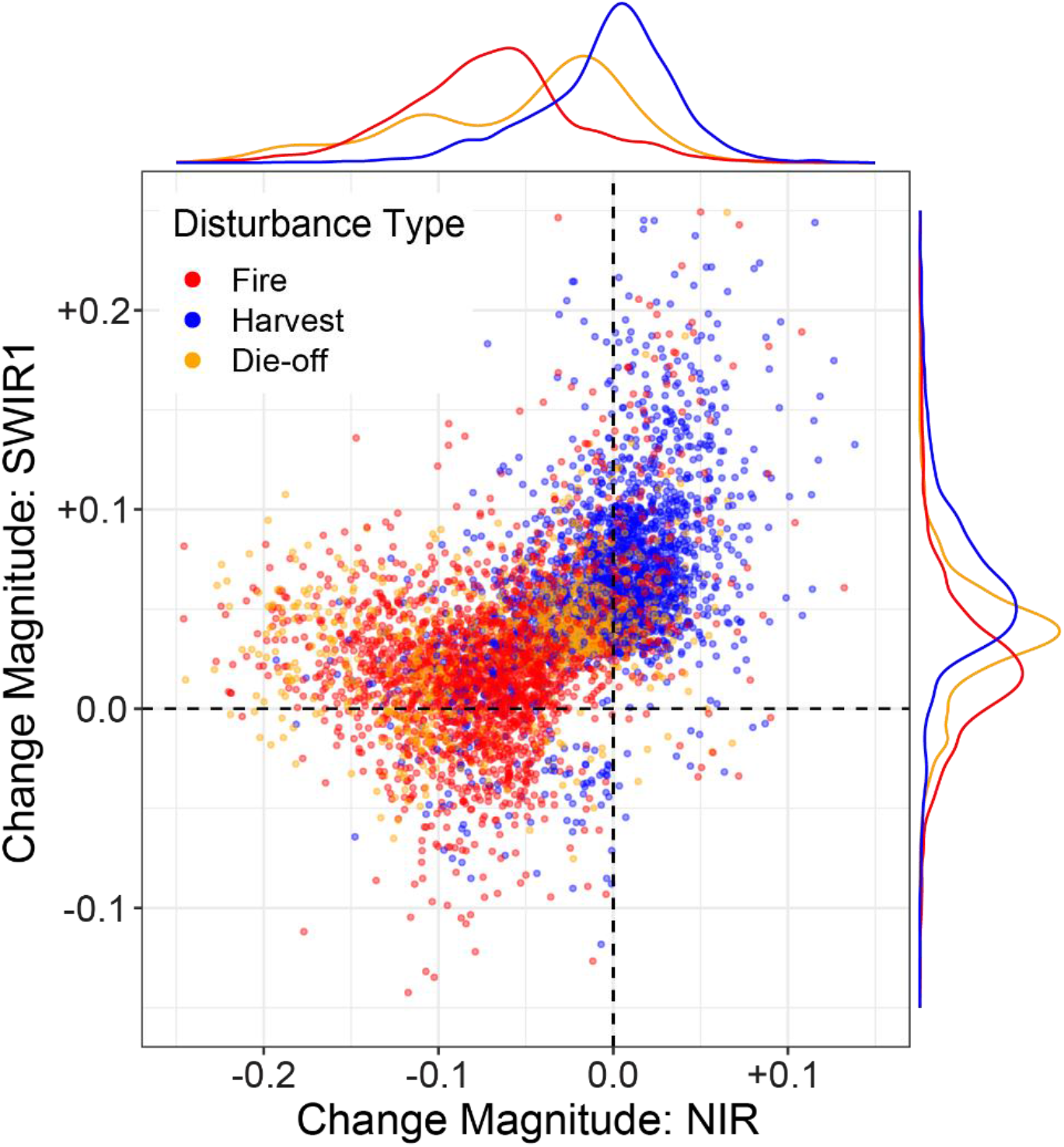
Scatterplot demonstrating the separability of different disturbance types in spectral change magnitude space. Shown is a sample of 25000 CCDC model breaks. Colors indicate the type of disturbance in the disturbance reference data set. The change magnitude on the x-axis indicate the change magnitude in near infrared reflectance (NIR, or Band 4 in Landsat TM), while the y-axis shows the change magnitude in the first shortwave-infrared band (SWIR1, or Band 5 in Landsat TM). Note that the actual model used all the spectral bands as inputs.

**Supplementary Figure S5.**
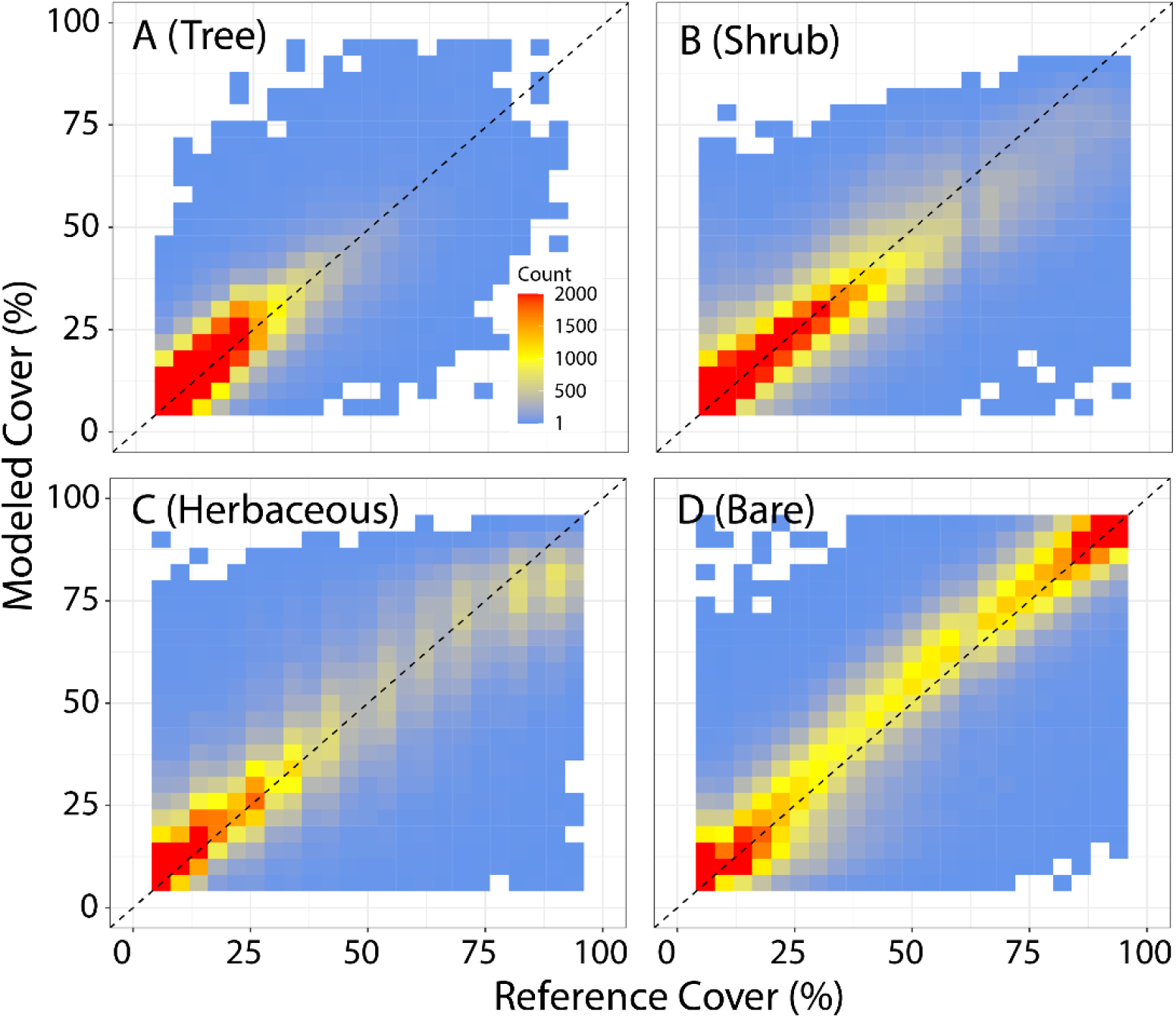
Model performance for prediction of fractional cover compared against out of sample reference data sampled from the NLCD tree cover and RCMAP data products. Model performance is shown for a) tree cover, b) shrub cover, c) herbaceous cover, and d) bare ground cover.

